# ATRAP - Accurate T cell Receptor Antigen Pairing through data-driven filtering of sequencing information from single-cells

**DOI:** 10.1101/2022.08.31.506001

**Authors:** Helle Rus Povlsen, Amalie Kai Bentzen, Mohammad Kadivar, Leon Eyrich Jessen, Sine Reker Hadrup, Morten Nielsen

## Abstract

Novel single-cell based technologies hold the promise of matching T cell receptor (TCR) sequences with their cognate peptide-MHC recognition motif in a high-throughput manner. Parallel capture of TCR transcripts and peptide-MHC is enabled through the use of reagents labeled with DNA barcodes. However, analysis and annotation of such single-cell sequencing (SCseq) data is challenged by dropout, random noise, and other technical artifacts that must be carefully handled in the downstream processing steps.

We here propose a rational, data-driven method termed ATRAP (Accurate T cell Receptor Antigen Paring) to deal with these challenges, filtering away likely artifacts, and enable the generation of large sets of TCR-pMHC sequence data with a high degree of specificity and sensitivity, thus outputting the most likely pMHC target per T cell. We have validated this approach across 10 different virus-specific T cell responses in 16 healthy donors. Across these samples we have identified up to 1494 high-confident TCR-pMHC pairs derived from 4135 single-cells.

## Introduction

T cells are essential for immune protection and play a critical role in the immune response to pathogens or cancer, where they directly kill infected or malignant host cells or orchestrate the response of other immune cells. Recognition is mediated through the heterodimeric T-cell receptor (TCR) expressed on the surface of T cells, which engages specifically with a peptide antigen (p) displayed in the MHC. Accurate specificity and broad coverage of antigen recognition is obtained through somatic recombination of the genetic loci, V(D)J, that encodes the α (VJ) and β (VDJ) chains of TCR. The process creates an extensively variable and dynamic repertoire, with an estimated 10^7^ distinct αβTCRs in an individual (Arstila et al., 1999; Davis & Bjorkman, 1988).

Due to this diversity, the individual TCR transcripts can be used as endogenous cellular barcodes inherited by the T cell progeny. This has been utilized for providing quantitative insight into TCR diversity (Robins et al., 2009), to trace lineage decisions of T cells (Gerlach et al., 2013) and to monitor the dynamics of T cells across immune-related diseases, such as infectious disease (Dziubianau et al., 2013; Hou et al., 2016), cancer (Kirsch et al., 2015; Sherwood, 2013; S. Q. Zhang et al., 2018) and autoimmunity (Acha-Orbea et al., 1988; Madi et al., 2014). Most of such TCR repertoire studies have been confined to bulk experiments, tracing the TCR β chain because of its greater diversity (compared to the alpha chain) and because it is less ambiguous due to allelic exclusion (Bergman, 1999). However, accurate pairing of the variable TCR α and β regions is valuable for uncovering the biological function of a T cell and is generally lost in bulk experiments since the transcripts are separately encoded. Moreover, we and others have earlier demonstrated such approaches are suboptimal for the characterization of TCR specificity, and that this characterization is dependent on both the a and b chains (Montemurro et al., 2021).

To accurately obtain TCR αβ-sequence-pair single-cell sequencing platforms can be applied to simultaneously capture both TCR chains, while retaining cell origin information. To further assign specificity information to such TCRs, T cells can be stained with barcode-labeled pMHC multimers to simultaneous identify pMHC specificity and TCR sequence of individual cells (Bentzen et al., 2016; S. Q. Zhang et al., 2018). Moreover, via DNA barcoded antibodies, the platform facilitates screening of surface proteins to distinguish cellular subtypes and enables cell hashing to trace origin of a given cell to e.g., a given donor, sample, or time-point, which is highly valuable in patient-studies.

We deployed the droplet-based single-cell platform from 10x Genomics. Ideally a droplet contains a single cell with all its analytes and a gel-bead in emulsion (GEM). The gel-bead contains barcoded primers which ensures tracing of transcripts back to the cell-of-origin, referred to as GEMs. While the platform is highly promising, the sequence deconvolution is associated with substantial noise, and challenging to discriminate true from false signals. Common confounding factors include stochastic gene expression, cell cycle variations, apoptosis, and technical artifacts such as multiplet capture, contamination, dropout, and batch effects. Dropout and stochastic gene expression both result in zero-inflated gene counts and are typically insensitive to low expression levels (Buettner et al., 2015; Kharchenko, Silberstein, & Scadden, 2014; Yamawaki et al., 2021). Multiplet capture is the event of capturing two or more cells in a single GEM and it is proportional to the capture rate of cells introduced to the system (Bloom, 2018; Zheng et al., 2017). The capture rate is determined by the rate of pulsing cells relative to the rate of gel-beads. Thus, to include even low frequency cell populations, the capture rate must be high at the expense of introducing more multiplets. Contamination is particularly an issue when including analytes such as pMHC multimers which may be dissolved in cell suspension (Gaublomme et al., 2019). The platform has no means of controlling how ambient analytes and their barcodes are partitioned with gel-beads in emulsion (GEMs) which leads to GEMs that appear like multiplets or consist of ambiguous annotations from multiple analyte barcodes. The reverse issue arises from the risk that analytes may dissociate from the cell before capture. The listed confounders may result in both false positive and false negative discoveries

The main concerns when screening for TCR specificity are nonspecific binding of pMHC and/or cell hashing analytes, incomplete TCR annotation, and T cell multiplets. Nonspecific binding and T cell multiplets may completely dilute the signal from actual interactions, while incomplete TCRs which are missing the annotation for either α- or β-chain render the single-cell setup superfluous. To ensure that a screening is fully exploited and interpreted correctly, we set out to develop a data driven algorithm that facilitates a consistent and reproducible TCR categorization (clonotyping), peptide-MHC (pMHC) annotation, and antibody-based cell hashing referencing of the donors and their HLA profile.

We applied this algorithm to two datasets, each derived from screening PBMCs from 16 healthy donors for T cell recognition against common viruses. In total, we evaluated TCR recognition against 10 different pMHC multimers, each labeled with their unique barcode. We demonstrate that following the filtering steps described here we can obtain a confident pairing of pMHC specificity and TCR sequence. This strategy will open novel opportunities to evaluate the structural interplay and the sequence-driven signatures of pMHC recognition at large scale.

## Results

### Parallel capture of TCRαβ sequences, peptide-MHC specificity and sample origin from single-cells

To obtain single-cell-derived triad information on TCR sequence, pMHC specificity, and sample origin; we stained peripheral blood mononuclear cells (PBMC) from a total of 16 different healthy donors (Table 1). All samples were stained with the same panel composed of 10 different viral-derived peptide-MHC (pMHC) multimers, each labeled with a unique barcode for that specificity and a common fluorescent label (allophycocyanin (APC)) (Fig. 1) (Table 2). To serve as an experimental control for the purity of the isolated T cells, we moreover stained the cells with three additional viral-derived pMHC multimers bearing a different fluorochrome (phycoerythrin (PE)) and labeled with their own unique DNA barcode (Supplementary Table 2). We sorted only the APC-labeled pMHC multimer binding T cells (and hence deselected the PE-labeled T cells) and included these in the down-stream single-cell processing.

**Figure 1.**
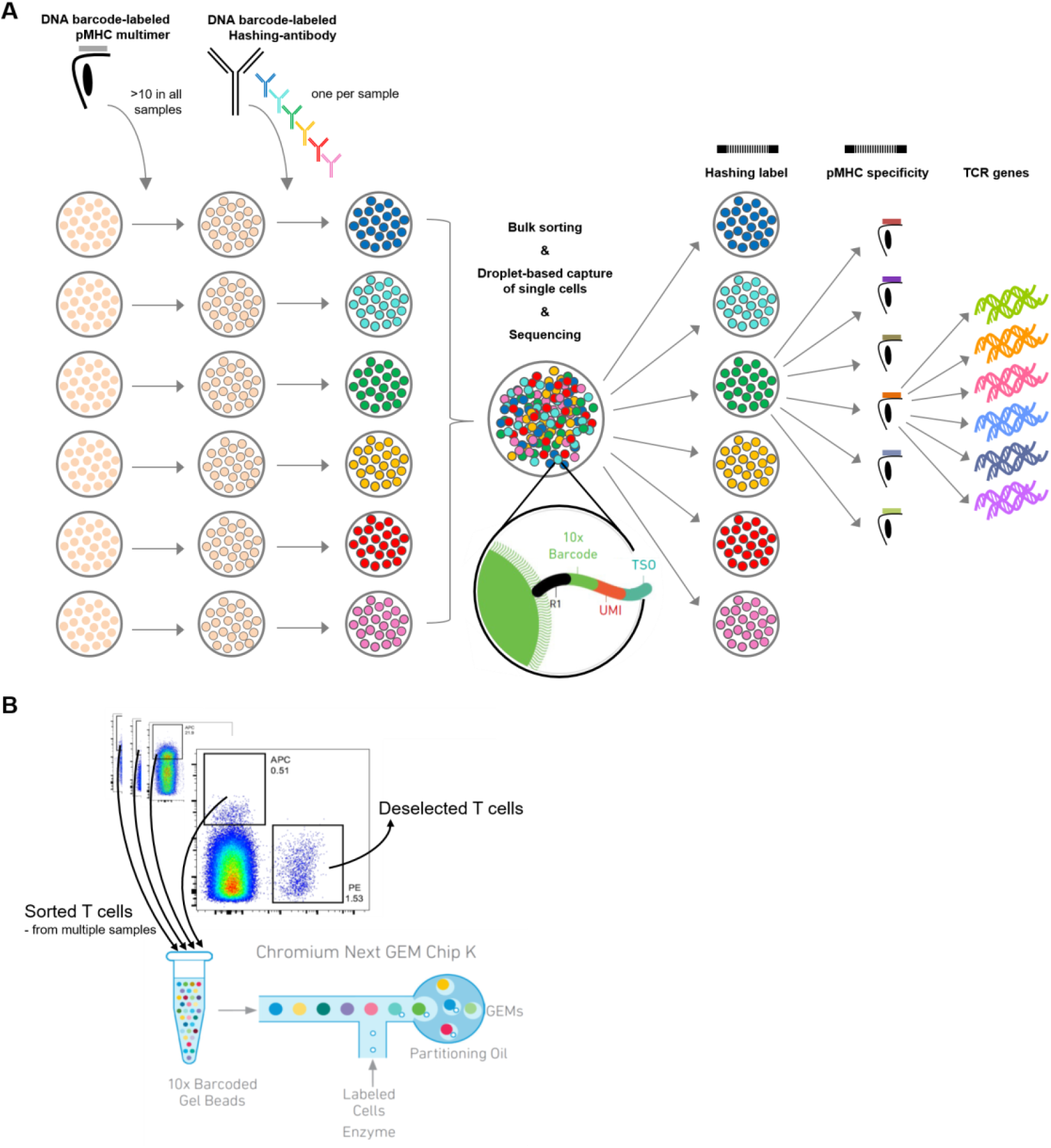
a) Schematic of the experimental strategy. All samples are incubated with the same library of barcode-labeled pMHC multimers and subsequently with a sample-specific barcode-labeled hashing antibody to individually label cells derived from a given sample. Multimer-binding cells from all samples are sorted in bulk and processed through the 10x Chromium workflow. The sequencing output simultaneously captures the sample barcode, the pMHC barcode and the TCR sequences, which are all matched to a single cell based on the 10x-barcode. This also provides the means of retrospectively assigning each cell to their sample of origin, via the sample specific hashing barcode. b) Example showing how the APC labeled pMHC multimers are sorted collectively from all samples into one tube that is further carried into the 10x workflow. The PE labeled pMHC multimers are not sorted and hence deselected. A total of 1800 APC labeled cells are sorted from each donor. Here showing BC126 (large dotplot) and BC341 (small dotplot).

Prior to sorting, each sample was stained with a distinct hashing antibody to provide a sample identification barcode associated with the GEMs of the resultant single-cell data set. This is done to enable mixing of cells from different samples, while retaining the information of sample origin, and utilizing the capacity of capturing 6,000-10,000 cells per lane in the 10xGenomics workflow. This is essential when capturing T cells based on their specificity since the MHC multimer positive population is generally of low frequency (<1% of CD8 T cells). When several samples are mixed in the process of running the single-cell analysis, all mRNA and DNA barcodes (derived from hashing antibodies or the MHC multimers) associated with a given cell will be encoded with the same 10x-barcode, proving the GEM association (Fig 1) (Supplementary Table 1).

### Total data from simultaneous capture of cell, TCR, pMHC and SampleID

The single-cell data is annotated using 10x Chromium Cellranger multi v6.1. This results in each GEM being quantified by a count of unique molecular identifiers (UMIs) (Kivioja et al., 2011) for the three components (TCR, pMHC and sample hashing) based on transcripts of TCR α- and β-chains, barcodes co-attached to pMHC multimers and barcodes co-attached to cell hashing antibodies (Supplementary Table 2).

To obtain the data presented here, a total of 1800 pMHC multimer positive cells were sorted per donor irrespective of the frequency or the number of different antigen-specific T cell responses in a given sample, accumulating to a total of 28,800 cells sorted. An estimated 45% of the sorted cells are lost in the process of loading on the Chromium, hence approximately 15,700 pMHC multimer labeled cells were included in the 10x experimental workflow. Initially, each GEM was annotated based on the most abundant transcripts from TCRαβ, pMHC, and cell hashing. However, this can lead to erroneous annotations, as the noise level can differ substantially for the different reagents, resulting in different levels of UMIs.

Based on raw, unfiltered data, we found 6,073 GEMs which contained all three components, i.e. TCR, pMHC and sample hashing, corresponding to 40% of the loaded cells (Fig. 2a). 716,069 GEMs only contained one or two of the components, with the majority containing only the cell hashing barcode (n=677,502) and the second largest share containing cell hashing as well as pMHC barcodes (n=37,277). This number vastly exceeds the number of cells in the assay (15,700 cells loaded) and indicates contamination from ambient barcodes in suspension. This is further supported by the observation that the sample hashing UMI count was significantly higher (p < 0.0005, Mann-Whitney U) in the 6,073 GEMs containing a TCR compared to the GEMs void of TCR (Fig. 2b). 43,455 GEMs captured a DNA barcode associated with the pMHC library and only 14% of these were completed with TCR transcripts and sample hashing barcodes. In the GEMs containing a TCR, 84% were completed with all three components, i.e. included hashing and pMHC barcodes, while less than 0.05% of these GEMs were void of both sample hashing and pMHC barcodes. In the following, we will only consider the 6,073 GEMs containing all three components, while taking into account that the high degree of noise also affects these seemingly completely mapped GEMs.

**Figure 2:**
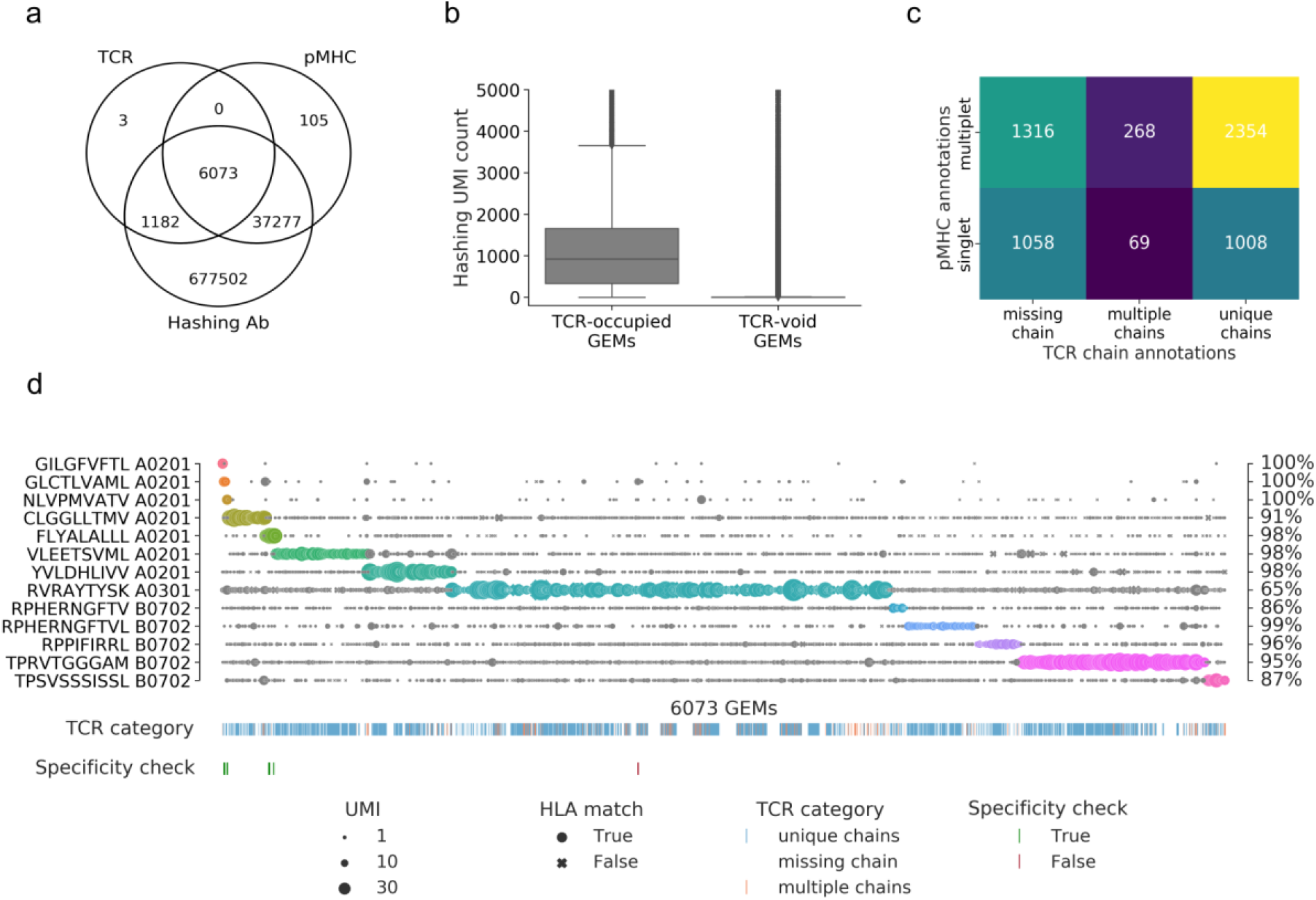
a) Venn-diagram of the content of all GEMs from 10x Chromium drop-seq. Each GEM is expected to contain three components: transcripts of TCR and DNA barcodes from the target pMHC multimer as well as the sample hashing antibody. The Venn-diagram illustrates the extent of GEMs with complete capture (capture of all three components) in contrast to the GEMs with incomplete capture (capture of a subset of components). b) Comparison of distributions of UMI counts of sample hashing barcode between GEMs that contain TCR transcripts (TCR-occupied GEMs) and GEMs that do not contain TCR transcripts (TCR-void GEMs) (p < 0.0005, Mann-Whitney U). c) Matrix of the distribution of pMHC singlets and multiplets across GEMs with TCRs either missing a chain, detected with multiple chains, or with a single, unique αβ-pair. The counts are given for each field and illustrated by a color. The lighter color represents higher counts. d) Scatterplot of all detected pMHC barcodes (y-axis) within each of the 6073 GEMs (x-axis) that contain all three components: TCR, pMHC and sample hashing. In each GEM the most abundant pMHC is marked by a color, while the remaining pMHCs in the GEM are gray. The marker size reports the UMI count of the given pMHC and the shape recounts whether the HLA allele of the pMHC matches the HLA haplotype of the donor, which is deduced from sample hashing. The fraction of HLA matches within the GEMs displaying a given specificity is annotated to the right of the plot. The first colorbar indicates the type of TCR chain annotation; whether the TCR has a unique αβ-pair, is missing a chain or consists of multiple chains. The second colorbar is a specificity check against the specificity databases IEDB and VDjdb. Colors highlight the GEMs where the CDR3αβ sequences are contained in the databases. The green color represents a match between the database pMHC and the detected pMHC, while red indicates a mismatch.

The GEMs are distributed across three categories of TCR and two categories of pMHC observations: GEMs either missing a TCR chain, contain multiple TCR chains, or contain a unique TCRαβ-pair and GEMs containing either a single or multiple pMHC barcodes (Fig. 2c). Sample hashing multiplets constitue 100% of GEMs containing sample hashing barcodes, and there is both a large proportion of pMHC multiplets (65%) and GEMs missing either α- or β TCR-chain (39%), hence, multiplets of pMHC and sample hashing is the predominant issue. Few GEMs were detected with multiple TCR α- or β-chains (6%). This may be caused partly by naturally occurring multiplets of α-chain (4%), due to the incomplete gene restriction of the thymocyte during negative selection (Elliott & Altmann, 1995; Petrie et al., 1993), or due to experimental features of the 10x platform causing an expected 6.9% of multiplets based on the number of cells loaded in our experiment.

Without further filtering, the pMHC-TCR pairing is subjected to extensive noise (Fig.2d) and we capture all the 10 DNA barcodes associated with the APC-labeled pMHCs in a varying number of GEMs. Importantly, the three negative control responses (GIL A0201, GLC A0201, and NLV A0201), which were present in the donors but not sorted, are only captured in a few GEMs. This indicates that the cell isolation via sorting is effective in terms of capturing only the desired cells and relevant pMHC-associated barcode-labels. The most frequently detected pMHC across all GEMs is RVR A0301, which is present with high UMI counts across all GEMs. Only RPH(10-mer) B0702-associated UMIs was consistently detected at low numbers per GEM. It was also evaluated whether the HLA allele of the pMHC matches the HLA haplotype of the donor(s) given via cell hashing (Fig. 2d). Typically, the mismatches are found in GEMs where the most abundant pMHC is detected at low UMI counts while the matches consist of GEMs with higher pMHC UMI counts. Of the 65% GEMs containing pMHC multiplets (Fig. 2c), 13% contained two or more pMHCs at the exact same UMI level (Supplementary Table 3), which may either represent noise or true cross-binding events.

The detected specificities in our data have been cross-referenced with the IEDB (Vita et al., 2019) and VDJ (Bagaev et al., 2020) databases (Fig. 2d). Based on the unfiltered data we found five TCR-pMHC matches (across 9 GEMs) and one TCR (1 GEM), which was annotated with a different pMHC (Fig. 2d). This latter is a case of a GEM with multiple pMHCs present with almost equal number of UMIs, where the most abundant pMHC is RVR A0301 (11 UMIs) and the second most abundant pMHC is GLC A0201 (9 UMIs), which is the peptide registered as target in IEDB and VDJdb.

The data in Fig. 2d suggests that most of the captured T cells interact with several of the screened pMHCs to a degree that exceeds the level expected from natural cross-recognition. Therefore, it is reasonable to assume that a large proportion of these multiplets are formed as a result of ambient pMHC leaking into GEMs.

### A data-driven filtering approach

From these observations, it is clear that a substantial part of the data consists of noise, i.e. GEMs with multiplets of pMHC and sample hashing, and that the data must be filtered for proper interpretation.

#### Clonotype annotation

The definition of T-cell clones (clonotypes) is fundamental for pairing a given TCR clonotype to its respective pMHC recognition. Initial clonotypes were called using 10x Genomics Cellranger which defines a clonotype as a set of cells that share identical receptor sequences at the nucleotide level, spanning the entirety of the V(D)J-C genes as well as the junction segments. Assuming reliable gene and CDR3 sequence calls by 10x Cellranger, we redefine clonotypes based on TCR annotation. Subsequently, GEMs with no clonotype annotation from 10x were annotated to existing clonotypes conditioned on matching VJαβ-genes and CDR3αβ sequences or as novel clonotypes. Similarly, clonotypes with identical VJ-CDR3αβ were merged to form larger groups of theoretically identical TCRs (Supplementary. fig 1). Merging GEMs of the same TCR is essential to make statistical inference based on those groupings e.g., determine expected pMHC target per clonotype. The outcome was a set of 2,441 TCR clonotypes across the 6,073 GEMs containing both TCR and pMHC. For the 337 GEMs containing TCR chain multiplets, the most abundant chain was for the subsequent analyses selected to represent the true TCR.

#### Defining pMHC recognition for selected TCR clonotypes

As we have seen earlier, not all GEMs within a given clonotype support the same pMHC target, and defining the pMHC target of a TCR based on individual GEMs thus results in contradicting annotations. The key to identify the expected target for a clonotype is therefore to determine which pMHC identity represents the majority of UMIs across all GEMs within a given clonotype. Fig. 3 illustrates an example from a pilot study which accentuates the importance of studying GEMs in ensemble rather than individually. Most GEMs are annotated with multiplets of pMHCs and across all GEMs the most abundant pMHC varies. While all pMHCs are found most abundant in at least one GEM, three pMHCs (TPR B0702, VTE A0101, and RAK B0801) are more often found most abundant (Fig. 3a). Although TPR B0702 is detected in fewer GEMs (136) than VTE A0101 (260) and RAK B0801 (186), TPR B0702 is present at generally higher UMI counts (Fig. 3b). It is evident that there is a difference in UMI distributions between the different pMHC within the GEMs of a given clonotype, and that TPR B0701 is the significantly most abundant pMHC across the ensemble of GEMs even though this pMHC is only present in a minority proportion of the GEM (Fig. 3b). Based on these observations, we argue that the significantly most abundant pMHC should be annotated as the expected binder for the given clonotype rather than annotating based on the majority.

**Figure 3:**
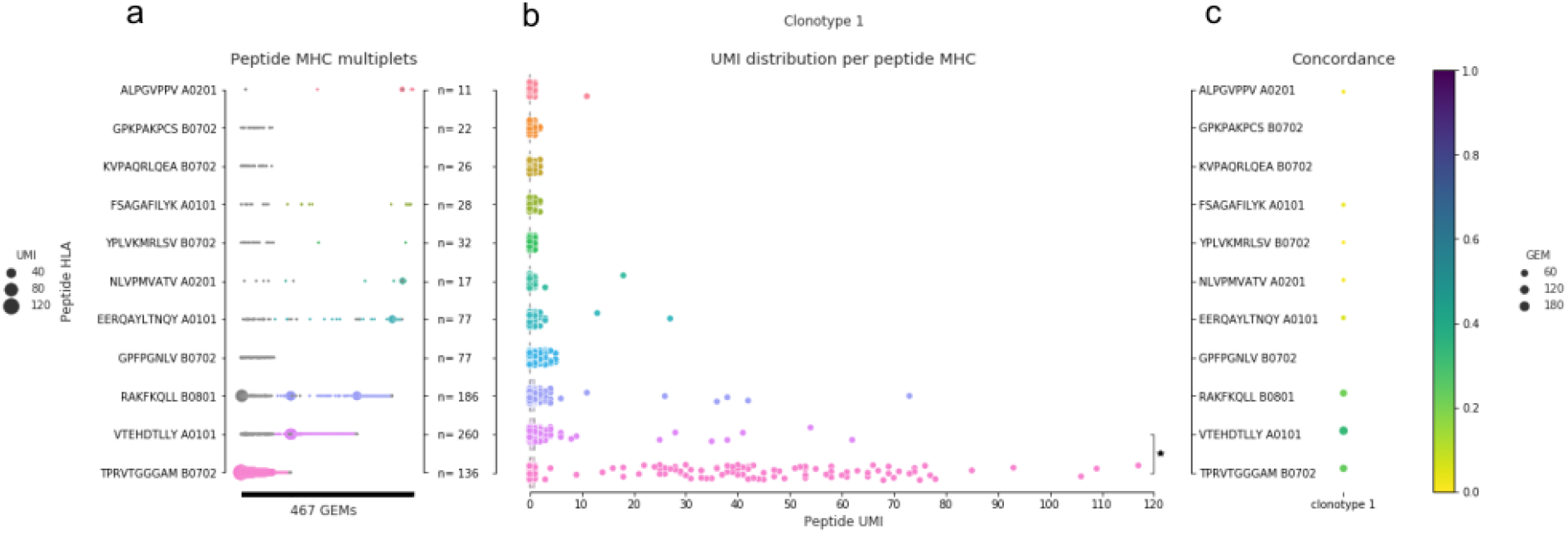
An example of pMHC concordance in clonotype 1 (example from pilot study). a) All detected pMHC (y-axis) in each GEM (x-axis, n=467) of clonotype 1. The marker size shows the UMI count for the particular pMHC in a given GEM, and the color indicates the pMHC with the highest UMI count, similar to what is shown in Fig. 1d. If two pMHCs are equally abundant in a GEM they are both colored. No marker means no detection of that pMHC in that given GEM. b) The compiled distribution of UMI counts for each peptide (assigning 0 UMI when the pMHC is not detected in a GEM). The asterisk marks that a Wilcoxon test showed that the UMI counts of TPR B0702 were on average higher than for VTE A0101 UMI counts. c) The specificity concordance across the GEMs of clonotype 1. Concordance is shown by a color gradient, i.e. the larger the fraction of GEMs supporting a given specificity the darker the color.

Having annotated the “true” pMHC of a given clonotype, one can next go back to the individual GEMs, and label GEMs where the most abundant pMHC corresponds to the expected binder, as “true”, and all others as “false”, and use these annotations to quantify the accuracy of the GEM annotations. Within each clonotype, one can compute a specificity concordance, i.e. the fraction of GEMs detected with a certain specificity (defined by most abundant pMHC, i.e. highest pMHC UMI per GEM) (Fig. 3c). In many cases across the full data set, the expected specificity for a clonotype coincides with the specificity, defined on a per-GEM level, resulting in high concordance. However, for some clonotypes e.g., clonotype 1, GEMs have diverging annotations and therefore lower concordance dispersed across multiple specificities (Fig 3). The clonotype visualized in Fig. 3 is specifically chosen to exemplify how this lower concordance can affect the analysis. For clonotype 1 the fraction of GEMs that support VTE A0101 (0.33) is higher than the fraction of GEMs that supports TPR B0702 (0.26). This results in an overall low concordance, and only by considering the complete ensemble of clonotype 1 GEMs, can the correct pMHC target be identified (Fig. 3b).

#### Improving concordance between GEM and clonotype annotation based on grid search on UMI features

To rationally filter data, an accuracy metric was defined, and optimized through the filtering process. For all specificities belonging to clonotypes with an assigned expected target, we calculated the overall accuracy as the proportion of GEMs where highest abundance pMHC annotation corresponds to the expected target of the clonotype. The raw unfiltered data yielded accuracy and average concordance scores of 69.6% and 83.8%, respectively. Next, we set out to investigate how different data driven UMI filters could improve these performance values, removing noise and artifacts from the data. This removal would also reduce the number of included observations, hence the performance of different thresholds for filtering the data was evaluated based on a tradeoff between increased accuracy and discarded number of GEMs.

We tested various thresholds on UMI count and UMI ratios, i.e. the ratio between the most abundant and second most abundant UMI feature, for pMHC and TCRαβ respectively. The optimal thresholds were chosen to maximize the weighted average between accuracy and fraction of retained GEMs to favor increase in accuracy above losing some GEMs. This filtering analysis resulted in optimal thresholds of 2 pMHC UMI counts and a ratio pMHC UMI counts between top one and two >1.The latter results in removal of GEMs where two pMHC were equally abundant for low UMI counts. The search did not result in thresholds imposing restrictions on neither TCR UMI counts nor TCR UMI ratio, which underpins that the TCRs with a missing chain as well as multiple chains also contribute to good performance. Imposing this filter yielded 5,061 GEMs (83% of total), 2,233 clonotypes (91% of total), and resulted in 96.4% accuracy, and a mean concordance of 93.6%.

#### Additional filters

Additional filters can be added to further clean the data. We compared how two filters, integrated in the 10x Genomics software, Cellranger, performed in removing potential noise from our data set (Supplementary Fig 2). The purpose of these filters is to evaluate, with high confidence, whether a GEM has captured a cell: “is cell” is defined based on the TCR transcript level in a given GEM and “is cell (GEX)” is defined based on the full transcript level (10xGenomics, n.d. a). Alternatively, viable cells are identified from the transcript data, independently of Cellranger, based on mitochondrial load and a minimum and maximum gene count per GEM. All three filterings are comparable (Supplementary Fig 2), and taken into account in the further evaluations. It is worth noting that, while the filterings based on the full transcript data might remove slightly more noise, the economic costs associated could propose that this should only be applied when the transcript data is required for additional purposes.

Cell hashing enables filtering based on sample demultiplexing methods such as Seurat hashtag oligo (HTO) demultiplexing to identify hashing singlets (Stoeckius et al., 2018) (Supplementary Fig 3 and Supplementary note). In this setup, cell hashing also enables filtering based on matching HLA between the donor haplotypes and the HLA of the detected pMHC. Additionally, depending on the subsequent use of the data, retaining only complete TCRs containing both α and β may be desirable. Including only GEMs where the TCR-pMHC pair is observed more than once, i.e. specificity multiplets, reduces the uncertainty described above. Below we investigate the impact of imposing such filters.

### Impacts of filtering

#### Evaluating filters by comparing TCR similarity across specificity

To objectively evaluate the performance impact of the presented filters, we define a quantitative evaluation based on the hypothesis that T cells binding the same pMHC (intra specificity) will share a higher sequence similarity compared to TCRs of different specificities (inter specificity) (Fig. 4). Thus, filtering away artifacts should increase intra-similarity while decreasing the inter-similarity. Here, the similarity score between two TCRs was calculated from the summed score of the pairwise α- and β-chain similarities calculated using a kernel method described in (Shen, Wong, Xiao, Guo, & Smale, 2012) and applied in (Chronister et al., 2021).

**Figure 4:**
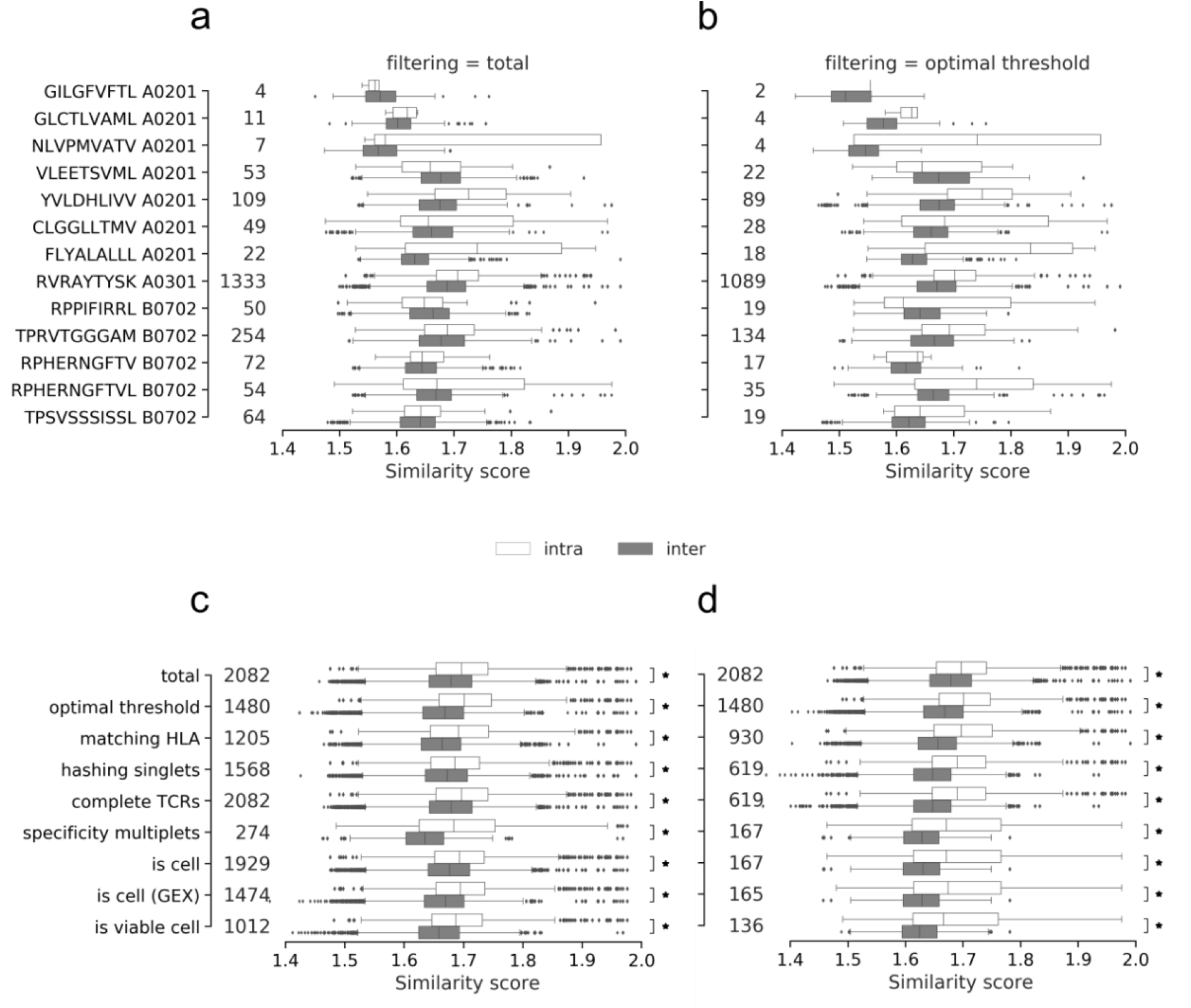
Intra- and inter TCR-similarity scores per peptide of the a) total (unfiltered) dataset, b) the data filtered by the optimized threshold. The similarity per peptide plots a) and b) illustrate the distribution of paired similarity scores for each clonotype (containing both α- and β-chain). For each pMHC each clonotype is compared to the remaining clonotypes of the same specificity (intra) and across specificities (inter). The count of compared clonotypes is listed just to the right of the y-axis in both a) and b). c) Displays the pooled intra- and inter-scores across all peptides for each of the filtering methods: total (no filtering), optimal threshold, matching HLA, hashing singlets, complete TCRs, specificity multiplets, “is cell” by cell-flagging, “is cell” by cell-flagging when including GEX data, and viable cell from analyzing GEX data. An asterisk marks filters where intra-similarity is significantly larger than inter-similarity (Wilcoxon, α=0.05). d) Displays the pooled intra- and inter-scores across all peptides for each of the filtering methods where each filtering is added cumulatively to the previously listed above it. An asterisk marks filters where intra-similarity is significantly larger than inter-similarity (Wilcoxon, α=0.05). The count of compared clonotypes is listed just to the right of the y-axis in both c) and d).

Based on this kernel similarity metric, the filters were tested individually and cumulatively, i.e. each filter was added to the previous set of filters. The general trend is that TCRs with the same specificity are more similar to each other than to TCRs of different specificities, when computing the intra and inter similarities per pMHC before and after filtering on the optimized UMI thresholds (Fig. 4a-b). Before filtering, nine out of 13 pMHCs displayed a higher mean intra-similarity than inter-similarity scores, whereas this number was 11 out 13 pMHCs when applying the UMI thresholds. The outliers before filtering were GIL A0201, VLE A0201, CLG A0201, and RPP B0702, while the outliers were reduced to VLE and RPP after filtering. Generally, the similarity scores often have a wide, overlapping range between the intra and inter categories. The three pMHCs that were deselected during sorting, GIL A0201, GLC A0201, and NLV A0201, are only detected in a few TCR binding events. To enhance the power of comparison, the intra and inter scores were pooled respectively across the individual pMHCs (Fig. 4c-d). The results demonstrate that intra-similarity is significantly higher than inter-similarity at each filtering step, both individually and combined. Moreover, we observe that the differences between intra- and inter-similarity appear to increase as filters are cumulatively added and fewer observations are left (Fig. 4d). Particularly, the median inter-similarity score is lowered, suggesting that the filtering steps predominantly removes false-positives.

#### Evaluating filters across selected performance metrics

To compare the effect of the filters, the similarity scores were converted to the performance metric: AUC (area under the receiver operating characteristic (ROC) curve). Here, intra specificity comparisons are regarded as true positive observations and inter specificity comparisons as true negatives. Based on these performance metric definitions, we quantify the effect of each filtering step (Fig. 5), and find that the highest accuracy and highest average concordance is obtained by filtering on the optimal threshold (95.3% and 90.6%), while the highest AUC is obtained from filtering on specificity singlets (70.5%) (Fig. 5a). Expectantly, the accuracy and average concordance increases when the filters are imposed cumulatively (Fig. 5b). The accumulation of filters also results in drastic reduction of the GEMs, and it is evident that one must carefully weigh out the need for specificity over sensitivity when selecting the desired set of filters.

**Figure 5:**
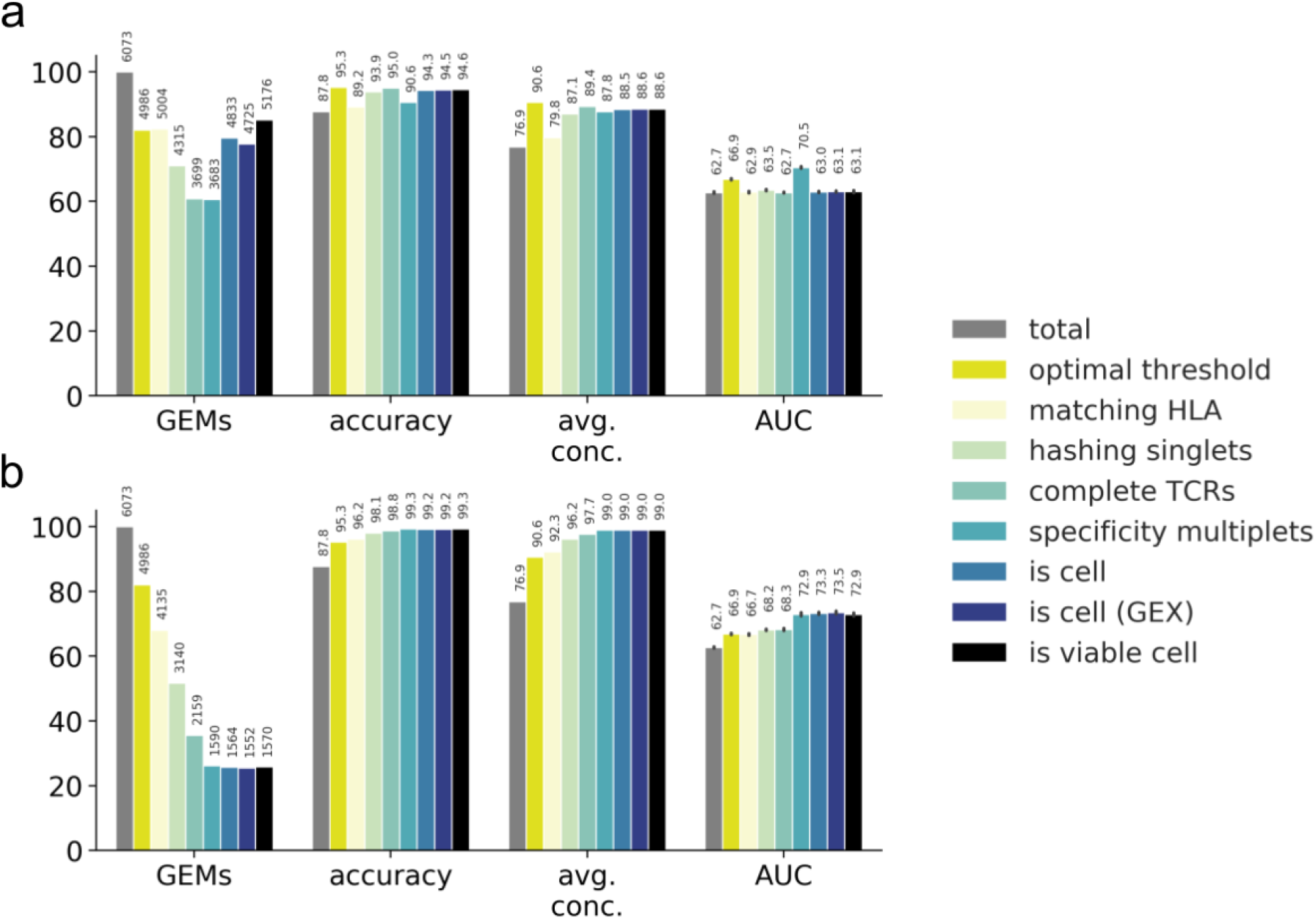
Performance metrics for evaluating the filtering steps. Performance is measured by number and ratio of retained GEMs (GEMs), accuracy defined by proportion of GEMs where most abundant pMHC matches the expected binder (accuracy), average binding concordance (avg. conc.) and AUC of similarity scores (AUC). The filtering steps consist of total (raw, unfiltered data), optimal threshold obtained from grid search, matching HLA, hashing singlets identified from Seurat HTO demultiplexing, complete TCRs with a unique set of α- and β-chain, specificity multiplets such that each TCR-pMHC pair must be observed in two or more GEMs, is cell defined by 10x Genomics Cellranger, is cell (GEX) defined by Cellranger where GEX data is included, and is viable cell defined by mitochondrial load and gene counts. a) Presentation of the individual effect of each filter. b) Presentation of the accumulated effects of the listed filters.

We conclude that the minimal filtering must include optimal threshold and matching HLA between pMHC and donor haplotype. Filtering on specificity multiplets would inherently result in more reliable observations, risking the removal of rare, low-avidity binding events. Generally, we did not find that including GEX data improved performance considerably. Finally, filtering on incomplete TCRs yields the second highest accuracy and average concordance. Unfortunately, the filter almost halves the number of GEMs. Hence, this filtering should be considered depending on future use of the data.

### Inspecting the filtered data

To determine the impact of the filtering steps, we have compiled the binding concordance for all clonotypes and applied three selected filtering steps: a) the raw, unfiltered data, b) filtering on optimal UMI thresholds and matching HLA, and c) additionally filtering on complete TCRs (Fig. 6). The raw, unfiltered data displays many clonotypes where the most abundant pMHC in GEMs of a given clonotype are dispersed across multiple of the screened pMHCs (Fig. 6a). When imposing the recommended set of filters, optimal threshold and HLA match, the outliers are greatly reduced, although not all low-concordance GEMs are removed (Fig. 6b). By additionally filtering on complete TCRs even fewer outliers are left (Fig. 6c). Note again that we have purposely deselected T cells specific for GIL A0201, GLC A0201, and NLV A0201, explaining the few observations for these otherwise frequently recognized epitopes.

**Figure 6:**
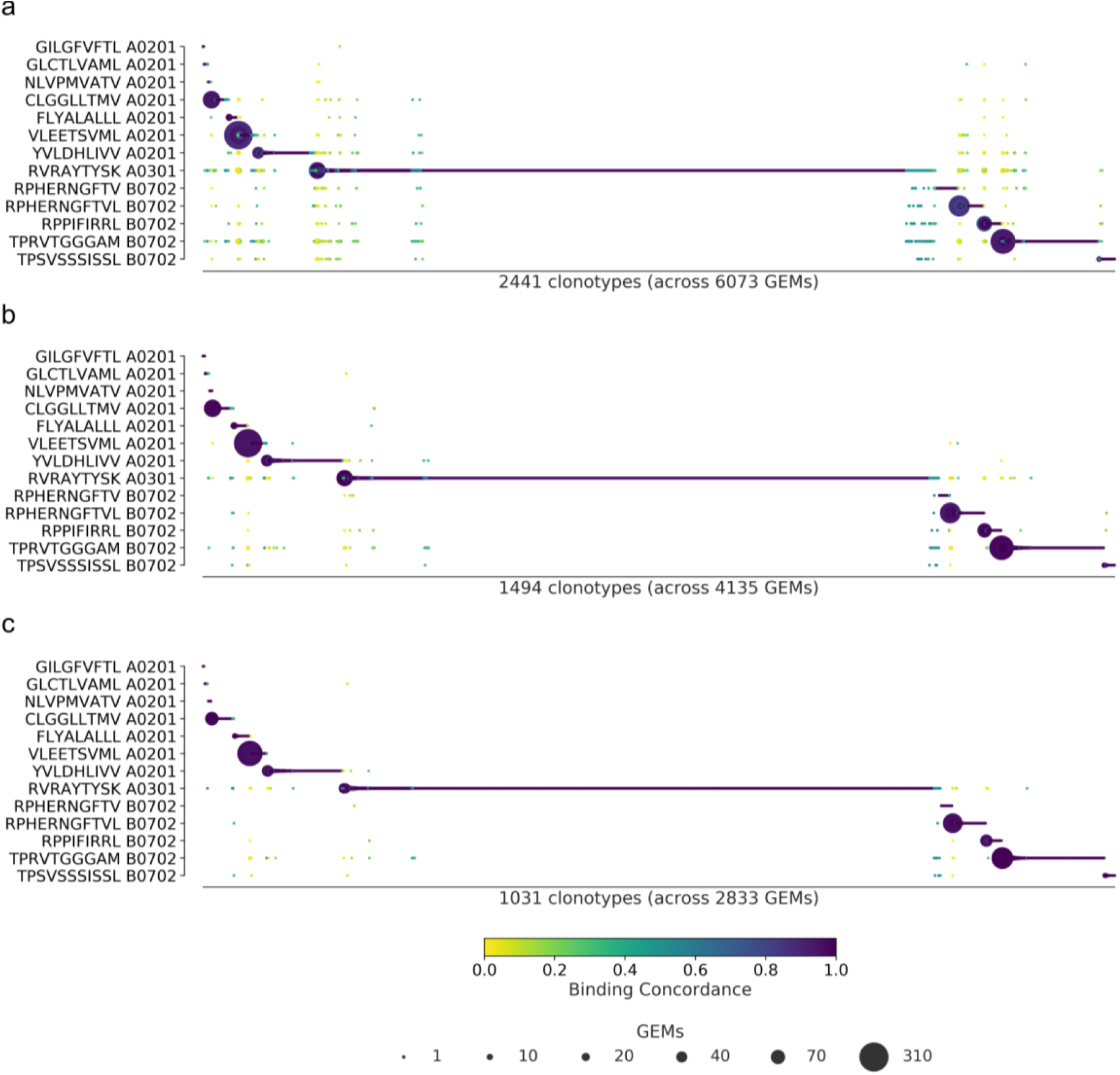
Specificity per clonotype. The library peptides are listed on the y-axis and each clonotype is represented on the x-axis. Below the x-axis is annotated the total number of clonotypes and GEMs in the presented data. The marker size shows the number of GEMs supporting a given specificity. The color indicates the binding concordance which is calculated as the fraction of GEMs within a clonotype that support a given pMHC. The higher the concordance, the larger the fraction of supporting GEMs. The three plots illustrate the impact of three filtering criteria. a) Presents raw data with no filtering applied. b) Presents data filtered on optimal threshold and HLA matches. c) Presents data filtered as in b) with the additionional requirement of only complete TCRs.

Many of the remaining low concordance GEMs still suggest the improbable event of cross-binding across HLA restriction. We suspect that these are artifacts that we have not successfully removed. When the most strict filtering is imposed (Fig. 6c) there are 56 GEMs (out of 2833) with a binding concordance of 0.5 or lower, which will be referred to as outliers. 50 of those GEMs contain pMHC multiplets. 94% of the multiplet outliers actually do contain the pMHC which defines the high-concordance GEMs, however, at a lower UMI count. In the GEMs with multiple pMHC annotations, the HLA is conserved across the pMHCs in 14% of the cases. In 68% of the cases the HLAs are different, but still match the HLA haplotype of the donor given by the cell hashing. Of the 56 outliers, the most dominant pMHCs are RVR A0301 (41%) and TPR B0702 (27%). Prior to filtering the data, six clonotypes were identified which were already registered in IEDB and VDJdb, five with matching pMHC and one with a different annotation than in our observation (Fig. 2d). The five matching clonotypes (9 GEMs) were successfully retained, while the mismatching clonotype (1 GEM) was filtered away.

### Comparing single-cell data with fluorescent-based pMHC multimer screening

#### Investigating dominant clones

Beyond mapping the landscape of known TCR-pMHC interactions, single-cell screening enables investigation of T cell repertoire diversity. The high resolution both reveals the specificity and the TCR clonality within the individual T cell populations, which is not possible to recover in classical stainings using fluorescent labeled pMHC multimers (fluorescent multimers). The T cell diversity in the nine donors towards the set of analyzed pMHCs reveals a clear hierarchy with dominant responses in fluorescent multimer staining (Fig. 7a), however the clonality of each specificity is only available via single-cell data (Fig. 7b). Here ATRAP represents data filtered by optimal UMI thresholds and matching HLA between pMHC and donor haplotype (given via cell hashing). Single-cell screening further enables comparison of the clonal distribution and the total clonal size per specificity. In this respect, the samples BC328 and BC62 are strikingly similar in their distribution of expanded clones. They both display a large and broad response towards RVR A0301 and two smaller responses towards RPP B0702 and TPS B0702 which are both dominated by a single clonotype. Further, most peptides elicit diverse relative responses between samples. For example, RPP B0702 is the dominant response in samples BC328 and BC62, but the minority response in sample BC314. Sample BC300 contains primarily small clones, i.e. fewer cells in each clonotype, however, this sample is generally represented with low amounts of total data (46 GEMs). Of note, small clones might be a result of suboptimal single-cell capture, or because high-frequency responses can potentially mask any lower frequency responses present in a given donor (Supplementary Table 4) when only 1800 cells are sorted from each sample. Samples represented with many GEMs are expected to be fully covered and therefore may contain more different expanded clonotypes, as sample BC360.

**Figure 7:**
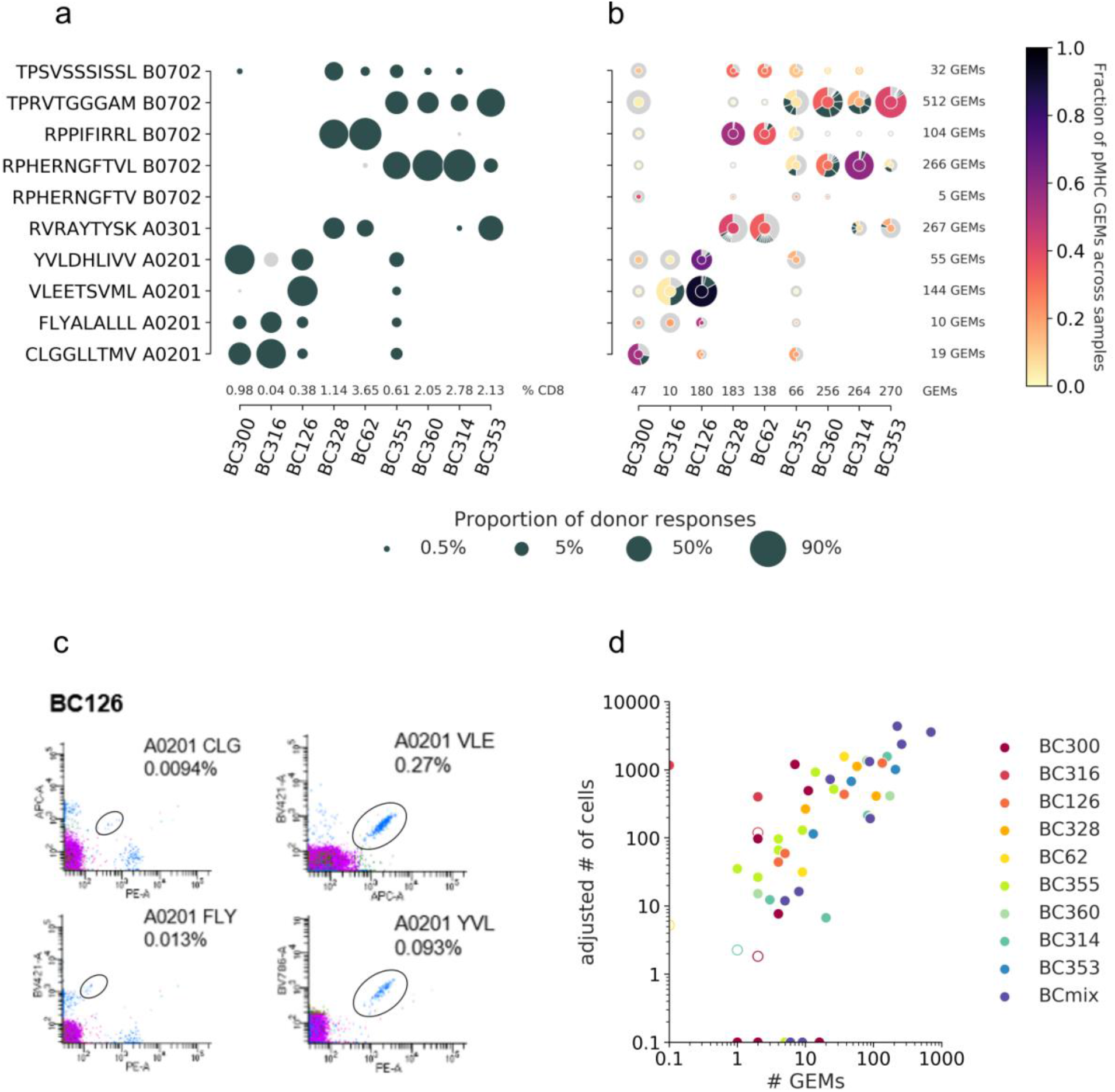
T cell diversity per peptide across the individual samples. The nine samples, PBMCs from nine individual donors are represented on the x-axis. The marker size defines the distribution of T cells recognizing a given peptide, normalized per sample. a) The T cell frequencies are visualized as the proportion of a given multimer positive response within a donor. The black markers represent responses detected above the threshold, i.e. ≥10 cells and ≥ 0.002% of total CD8 T cells, or ≤10 cells but ≥0.01% of total CD8 T cells. The gray dots represent detected specificities below threshold but represented by ≥2 cells. Summed frequencies of detected responses within a donor are given as % of total CD8 T cells and listed just above the x-axis. b) The T cell frequencies are based on GEM counts normalized per sample from the single-cell data. Absolute GEM counts per sample are listed above the x-axis. The marker is colored by the fraction of GEMs within a specificity that originate from a given sample. Absolute GEM counts per peptide are listed to the right of the plot. The marker contains a donut diagram illustrating the distribution of clonotypes specific for the given peptide in the given sample. The wedge that represents the dominant clone is colored according to the center of the donut. Remaining clones (>1 GEM) are anthracite gray and all clonotypes only supported by one GEM only are pooled and represented by a single light gray wedge. Comparing the sizes of the T cell populations for each specificity per donor between the two screening methods in a) and b) yielded the following Spearman correlations: BC126 (1.00, p<0.0005), BC328 (0.90, p=0.006), BC355 (0.74, p=0.02), BC360 (0.89, p=0.04), BC314 (0.90, p=0.04), and BC353 (1.00, p<0.0005). c) Representative example showing the four different responses detected with fluorescent-labeled pMHC multimers in donor BC126. d) Correlation between T cell responses detected by fluorescent labeled MHC multimers (y-axis) and single-cell capturing (x-axis). Correlation is given by Pearson correlation coefficient 0.73 (p<0.0005). The responses from fluorescent-based screening is given as an adjusted number of cells based on the detected response frequency out of 1800 cells (see calculations in supplementary table 4). The hollow markers represent responses below detection threshold as described in a). The responses are colored by the donor-of-origin. BC mix corresponds to BC311, BC11, BC83, BC88, BC341, BC342, and BC76.

#### Evaluating ATRAP by “ground truth” of fluorescent-based pMHC multimer screening

The net-overlap of identified T cell responses between the two screenings (Fig.7a+b) is estimated to 0.63 by Matthew’s Correlation Coefficient (MCC). Most of the T cell populations detected by fluorescent multimers are also captured in the single-cell screening, reflected by a recall of 0.95. However, the single-cell capture of small T cell clones (Fig 7b) that were not detected using fluorescent multimers (Fig 7a), negatively impacts the precision, yielding a score of 0.71. These “false positives” could result from low affinity T cell clones where the fluorescent signal would not be distinguishable from the background. Importantly, most of these responses were only represented by 1 GEM per clonotype, (demonstrated by a light gray outer circle in Fig 7b). In only two cases T cell populations were detected with fluorescent multimers but not captured in single cells: BC316/CLG A0201 and BC62/ RPH(10mer) B0702 (Fig. 7a). The large T cell population of BC316/CLG A0201 was likely a technical artifact related to the barcode-labeled pMHC multimers.

We calculated the number of antigen specific T cells sorted per donor, based on the total number of sorted cells/donor (n=1800) and the frequency of each T cell population (Fig 7c and Supplementary Table 4). This number of sorted cells for a given specificity was strongly correlated with the numbers of single-cell GEMs assigned to the same specificity (Pearson correlation coefficient, PCC=0.73, p<0.0005). We also fitted a linear regression for T cell populations sorted and assigned with at least one adjusted cell count or GEM in the log-log space (R2=0.56). The regression indicates that ∼10% of sorted cells will be captured in a single-cell screening with TCR-pMHC information yielded by ATRAP.

## Discussion

Here, we have described and validated, ATRAP; a data-driven approach for Accurate Pairing of T cell Receptor and Antigen. We have successfully filtered single-cell 10x Genomics data to identify reliable TCR-pMHC interactions of up to 1494 clonotypes. The method can be adapted to any single-cell immune profiling data set and is highly transparent in the steps taken, allowing the user to choose appropriate stringency of filtering.

Our recommended approach of cleaning data with minimal elimination of GEMs is obtained by two sets of filters: 1) the optimized data-driven UMI thresholds combined with 2) information on matching HLA specificity (as obtained from donor-specific hashing). Increasing filtering is naturally at the expense of the number of GEMs which might reflect the trade-off between specificity and sensitivity of the assay. However, any benchmarking or validation is made difficult without a golden standard. Our best attempt at quantifying the impact of filtering is based on three metrics: annotation accuracy, binding concordance, and AUC of clonotype similarity for which ATRAP yielded the scores 96.2, 92.3, and 66.7, respectively. Evaluation of ATRAP with responses from fluorescent pMHC multimer staining revealed strong correlation (PCC=0.73, MCC=0.63) between the number of sorted T cells and the number of detected GEMs across all specificities.

Accuracy of pMHC annotation was based on selected clonotypes where the expected target was statistically distinct and UMI thresholds were set to optimize the annotation accuracy. Rare clonotypes are not considered in this metric and clones are not expected to display cross-reactivity amongst the included pMHC multimers. The optimal UMI thresholds are intended to remove observations deviating from the expected target. The identified UMI thresholds are data specific and cannot be universally applied, but must be fitted for individual experiments. The thresholds are based on the assumption that contamination will predominantly exist at lower UMI counts than actual binding events. This limits the sensitivity of the method in cases of low-affinity low-frequency interactions which otherwise might be of great scientific and clinical interest.

The binding concordance is a metric that highlights cross-reactive clonotypes. In assays where cross-reactivity is not an expected outcome, binding concordance can be useful to evaluate the clonotypes where an expected target could not be identified. On the contrary, for data where T cell cross-recognition is of particular interest, the binding concordance can be used to establish the relative TCR binding contribution of each of the attributed pMHC targets. Growing evidence point to the relevance of T cell cross-recognition in both infectious disease (Dowell et al., 2021) and cancer (Fluckiger et al., 2020). Hence, novel tools to interrogate this phenomena on a single T cell level is highly warranted.

The last evaluation metric, AUC of clonotype similarity, is based on the assumption that T cells sharing specificity have more similar TCR sequences than T cells of different specificities (Chronister et al., 2021). This approach showed increasing separation of intra specificities and inter specificities as filters were cumulatively added, indicating that non-specific binders were effectively removed. To further increase the AUC, discarding clonotype singlets (i.e. TCR clonotypes represented by only 1 GEM) was the best single filtering step to improve the AUC of similarity scores (AUC=70.5, Fig 5a). This likely reflects that a fraction of such clonotype singlets represents non-specific binding events. However, removing these as a standard procedure of ATRAP, results in a substantial loss of TCR capture, represented by all T cell specificities with a light gray outer circle in Fig 7b. Thus, when aiming for capture of very low-frequency T cell specificities, a balance should be made between including this more stringent filtering step, or including such events, as demonstrated here.

To the best of our knowledge only one other method (ICON) has been proposed to clean TCR-pMHC single-cell data (W. Zhang et al., 2021). ICON was developed based on the public 10x Genomics data which includes six negative control pMHCs (Boutet et al., 2019), and 44 pMHC for positive selection of T cell populations. Comparing our method with ICON suggests that we present a more flexible and customizable approach. Where ICON retained ∼30% of their original data (W. Zhang et al., 2021), the ATRAP method presented here allows varying yields, depending on the level of filtering applied. The optimal filtering combination of UMI thresholds and HLA matching retained ∼70% of the data, while the combination of all presented filters retained ∼26%. As ICON does not consider the donor haplotype information, ∼15% of their specificities contained HLA mismatches, and a number of the T cell annotations includes TCR chains of diverting clonotype definitions given by 10x Genomics. For both methods, a particular awareness should be assigned to properly handle the range of avidity displayed by different clonotypes. One clonotype may display natural low avidity towards its cognate target which might appear like noise in the comparison to other high avidity clonotypes. This diversity in signals is challenging to handle in a one-fit-all filtering process, and for projects with specific interest in low-avidity cell interactions, a specific focus should be addressed not to lose such information.

Effective pairing of TCR and pMHC will open new avenues to interrogate T cell recognition and the role of different T cell populations in pathogenic processes. Intensive efforts have been made to identify antigen specificity based on the TCR sequence (Gielis et al., 2019; Montemurro et al., 2021; Moris et al., 2021; Sidhom, Larman, Pardoll, & Baras, 2021; Weber, Born, & Rodriguez Martínez, 2021; W. Zhang et al., 2021), and access to both TCR α- and β-chain is important to improve such prediction strategies (Montemurro et al., 2021).

The coveted data is ensured via the ATRAP framework for single-cell data of TCRs and associated barcodes. The perspectives of further exploiting the transcriptomic information, allowing in-depth tracking of specific T cell subsets based on the clonotypes, suggests that we are on the verge of achieving substantial novel insight to T cell involvement and behavior in health and disease.

## Acknowledgments

We would like to thank all healthy donors contributing material to this study. This research was funded in part through the Independent Research Fund Denmark (DFF 7014-00055 to S.R.H. and M.N.), the Lundbeck Foundation (R322-2019-2445 and R324-2019-1671 to A.K.B, and R190-2014-4178 to S.R.H), the European Research Council, StG 677268 NextDART to S.R.H., and National Institute of Allergy and Infectious Diseases (NIAID), under award number 75N93019C00001 to H.R.P and M.N.

## AUTHOR CONTRIBUTIONS

H.R.P and A.K.B. conceived the idea, designed experiments, analyzed data, made figures and wrote the manuscript; A.K.B. and M.K performed experiments, H.R.P. and L.E.J. developed bioinformatic processing; S.R.H. and M.N. conceived the idea, supervised the study, designed experiments and analyzed data; S.R.H., M.N., M.K. and L.E.J. revised the manuscript.

## Methods

### Ethical approval

All healthy donor material was collected under approval by the Scientific Ethics Committee of the Capital Region, Denmark, and written informed consent was obtained according to the Declaration of Helsinki.

### Cell samples

Peripheral blood mononuclear cells (PBMCs) from healthy donors were isolated from whole blood by density centrifugation on Lymphoprep (Axis-Shield PoC) and cryopreserved at −150 °C in FCS (Gibco) + 10% DMSO.

### DNA barcodes and dextran conjugation

Oligonucleotides modified with a 5′ biotin tag were purchased from LCG Biosearch Technologies (Denmark). Read from 5’ to 3’, the oligonucleotides were designed with the 10x equivalent Read2N sequence, a 10 nt unique molecular identifier (UMI), a distinct 15mer nucleotide sequences (extracted from (Xu et al., 2009), a 9 nt UMI and ending in a 13 nt capture sequence complementary to the TSO of the 10x 5’ capture oligo (sequences are listed in Supplementary Table 1). Barcodes were dissolved to 100 μM in RNAse free water and stored at −20 °C. For a working solution the DNA barcodes were further diluted in PBS + 0.5% BSA + 1 mg/mL herring DNA + 2 mM EDTA to 2.17 μM and attached to PE- or APC- and streptavidin-conjugated dextran from FINA Biosolutions LCC (USA). The amount of DNA barcode attached to each new lot of dextran was titrated as described in Bentzen et al., 2016. DNA barcodes were attached by mixing with dextran-conjugate, followed by incubation, 30 min at 4 °C. DNA barcode-assembled dextran-conjugates were stored for up to 24 hours at 4 °C.

### Peptides and MHC monomer production

Peptides were purchased from Pepscan (Pepscan Presto) and dissolved to 10 mM in DMSO. UV-sensitive ligands were synthesized as previously described (Bakker et al., 2008; Rodenko et al., 2006; Toebes et al., 2006). Recombinant HLA-A*0201, HLA-A*0301 and HLA-B*0702, heavy chains and human β2 microglobulin light chain were produced in Escherichia coli. HLA heavy and light chains were refolded with UV-sensitive ligands and purified as described in (Hadrup et al., 2009). Specific peptide-MHC complexes were generated by UV-mediated peptide MHC exchange (Chang et al., 2013; Frøsig et al., 2015; Rodenko et al., 2006; Toebes et al., 2006).

### Generation of DNA barcode-labeled MHC multimer libraries

Unoccupied SA-binding sites on the DNA barcode-assembled dextran conjugates were used for the co-attachment of biotinylated pMHC molecules. 0.8 pmol pMHC monomer was mixed with 160 × 10^−15^ mol DNA-barcoded dextran-conjugate and incubated 30 min at RT. MHC multimers were diluted in PBS with 5.2 μM d-biotin (Avidity, Bio200) to 909 nM and incubated 20 min on ice. DNA-barcoded MHC multimers were stored for up 1 week at −20 °C (PBS supplemented with glycerol and BSA, final concentrations 5% and 0.5%, respectively). Immediately before staining barcode-labeled MHC multimers were thawed at 4 °C, centrifuged (5 min at 3,300g), and pooled (0.8 pmol of each pMHC/sample) to enable the detection of multiple T-cell responses in parallel. The pooled MHC multimers were centrifuged once more; 5 min at 3,300g, to sediment aggregates before the volume of the reagent pool was reduced by ultrafiltration to obtain a final volume of ∼80 μL of MHC multimers as described in Bentzen et al., 2016. Any aggregates in the MHC multimer reagent pool were sedimented by centrifugation, 5 min at 3,300g before addition to the cell sample.

### MHC multimer staining

Cryopreserved PBMCs were thawed and washed by sedimentation, 5 min, 390*g*, 4 °C, in RPMI + 10% FCS. Cells were further washed in a barcode-cytometry buffer (PBS + 0.5% BSA). 5 × 10^6^ cells were incubated, 60 min, 4 °C, with pooled DNA-barcoded multimers in a total volume of 100 μL (final concentration of each distinct pMHC, 8 nM), and washed three times by sedimentation, 5 min, 390*g*, 4 °C. 5 µl of Human TruStain FcX™ Fc Blocking reagent was added to a total of 50 µl cell suspension, and incubated 10 min, 4 °C. Hashing antibodies (Biolegend, TotalSeq™-C0251 anti-human Hashtag 1-10 Antibodies) were centrifuged 10 min, 14,000 x *g*, 4 °C, and 0.5 µl were added to each a distinct sample (Supplementary table 2), and incubated 15 min, 4 °C. Next a 5× antibody mix composed of CD8-BV480 (BD 566121, clone RPA-T8) (final dilution 1/50), dump channel antibodies: CD4-FITC (BD 345768) (final dilution 1/80), CD14-FITC (BD 345784) (final dilution 1/32), CD19-FITC (BD 345776) (final dilution 1/16), CD40-FITC (Serotech MCA1590F) (final dilution 1/40), CD16-FITC (BD 335035) (final dilution 1/64) and a dead cell marker (LIVE/DEAD Fixable Near-IR; Invitrogen L10119) (final dilution 1/1000) was mixed for each sample. The antibody mix was added to cell samples and incubated 30 min, 4 °C. Cells were washed three times in barcode-cytometry buffer and kept on ice until acquisition.

### Cell sorting

Cells were sorted on a FACS Melody (BD) into tubes containing 100 μL of PBS + 0.5% BSA (tubes were saturated with PBS + 2% BSA in advance). Using FACS Chorus software, we gated on single, live, CD8-positive and ‘dump’ (CD4, 14, 16, 19 and 40)-negative lymphocytes and sorted only APC-positive (PE-negative) cells within this population (Supplementary Fig 4 for gating strategy). Cells sorted from individual samples were collected into the same tube (Fig 1b). The sorted cells were centrifuged for 10 min at 390*g* and the buffer was removed.

### DNA barcode-labeled MHC multimer stained cells on 10x platform

We utilize the 10x 5’ v2 chemistry that allows the cell barcode to be appended at the 5’-end of transcripts, which is essential for capturing the CDR3 region of the V(D)J transcripts. In the 5’ chemistry, the template switch oligo (TSO) is encoded with a cell barcode, i.e. one unique 10x barcode for every Gel Bead-in-emulsion (GEM). The TSO thus comprises the capture oligo, whereas the poly-dT primer is added in free suspension. Reverse transcription is initiated from binding of the poly-dT primer at the 3’-end, and mRNA is captured when the reverse transcriptase enzyme switches at the 5’-end of the transcript to the TSO. All DNA barcodes, partially complementary to the 10x Genomics 5’ TSO, are captured directly onto the GEMs. Annealing and extension during the reverse-transcription reaction associates the cell barcode and unique molecular identifier (UMI) from the gel bead oligo with the pMHC and hashing antibody tags in parallel with the mRNAs in the same droplet.

Downstream processing of mRNA and DNA barcodes are performed according to manufacturer’s instructions (Chromium Next GEM Single Cell 5’ Reagent Kits v2 (Dual Index), with the Feature Barcode technology for Cell Surface Protein & Immune Receptor Mapping) (10x Genomics, USA). ∼15,700 cells were loaded (based on 55% recovery from 28,800 sorted cells) to yield a maximum of 9,000 cells with an intermediate/high doublet rate (6,9%). Targeted amplification was performed for 13 cycles and the products were separated according to size into <400 bp (DNA barcode-tags) and >400 bp (the TCR sequences) using 0.6x SPRIselect beads (Beckman Coulter, B23318). Separate processing of the >400 bp bead-bound TCR sequences and the <400 bp in solution DNA barcodes was conducted according to manufacturer’s instruction and the TCR amplification products were sequenced on a NovaSeq running a 150 paired-end program. DNA barcodes, TCR sequences and mRNA was sequenced to a depth of 13,332, 12,503, and 18,398 mean reads per cell, respectively.

## Bioinformatics

### Processing of 10x single-cell data

Hashing barcode reads, peptide-MHC barcode reads, and T cell gene expression reads, were provided in fastq format and were processed using 10x Genomics Cellranger multi v6.1.0 (10xGenomics, n.d. b). The relevant outputs were the unfiltered count matrices of DNA barcodes and gene expression as well as clonotype annotations of each sequencing contig containing CDR3α/β sequences, V(D)J-C genes and unique molecular identifier (UMI) counts.

### Postprocessing 10x Cellranger clonotyping

The raw contig annotations from Cellranger were selected for downstream analysis with filtering on incomplete and unproductive receptor transcripts. Incomplete contigs are not full length, i.e. do not span the V-gene start codon until the J-gene stop codon. Unproductive contigs contain a frameshift which either induces an early stop codon or completely removes the stop codon.

Clonotypes defined by 10x were merged when consisting of identical VJ-CDR3αβ, thus reducing functional duplicates.

Cellranger flags rare nucleotide transcripts as likely artifacts, meaning the GEMs are flagged as unlikely to contain a cell and are therefore not assigned a clonotype (10xGenomics, n.d. a). Therefore, GEMs that were not annotated with a clonotype were imputed by searching the duplicate-reduced clonotype set. If no match, a new clonotype ID was annotated to the GEM.

### Filtering based on gene expression

Filtering on gene expression data was performed as described in (W. Zhang et al., 2021). Low-quality GEMs such as doublets may be removed by excluding GEMs with more than 2500 genes. Dead cells may be removed by excluding GEMs with fewer than 200 genes and a ratio of mitochondrial gene expression to the total gene expression above 0.2.

### Demultiplexing samples via cell hashing

Cell Hashing uses oligo-tagged antibodies against ubiquitously expressed surface proteins to place a sample barcode on each single cell, enabling different samples to be multiplexed together and run in a single experiment. To demultiplex the samples the method presented by Stoeckius et. al was implemented (Stoeckius et al., 2018). The method clusters the normalized count matrix using k-medoid clustering into k clusters, *k = n*_*samples*_ + 1. For each barcode a negative binomial distribution is fitted to the pool of all clusters except the cluster with the highest average expression for the given barcode. Each GEM is classified as positive if the barcode value exceeds a 0.99 quantile threshold for the negative distribution, and otherwise classified as negative. If GEMs contain multiple barcodes which pass the threshold, the GEM is annotated as a doublet (Stoeckius et al., 2018).

### Defining the expected binder

The pMHC and cell hashing barcode annotations were merged with the T cell annotations on the GEMs which contained both TCR and pMHC attributes. Each clonotype is expected to have a preferred target within the pMHC library, thus each clonotype is evaluated to find the pMHC which is most likely to be that target. Each clonotype is evaluated to identify the expected target within the pMHC library. The pMHCs that are detected within the GEMs annotated to a given clonotype are compared by their UMI count distribution. The two pMHCs that have the highest mean UMI count are compared, testing the hypothesis that the expected binder will have a significantly higher mean UMI count than the other pMHC (Wilcoxon, α=0.05). Clonotypes of less than 10 GEMs were not tested. The clonotypes where the mean UMI of the top two pMHCs was significantly different were collected as a training set. The pMHC which had significantly higher mean UMI was annotated as the expected target and specificity annotations of the GEMs were individually evaluated. The GEMs in the training set where the most abundant pMHC matched the expected target were labeled as true interactions, and the rest were labeled as false interactions.

### Defining specificity concordance

Concordance is an indirect measure of how cross-reactive a certain clonotype is. Specificity concordance is defined as the ratio of GEMs of a single clonotype which are annotated to bind a particular pMHC. The more GEMs in a clonotype annotated to the same pMHC the larger concordance. If a clonotype is only detected with one pMHC the specificity concordance is 1.

### Grid search on UMI features

Based on the labels of the training set a performance metric, o, was defined to evaluate the accuracy at increasing thresholds for UMI count and UMI ratio of pMHC, α-chain, and β-chain. The UMI ratio measures multiplets and is defined as the ratio between the highest UMI count and the second highest UMI count in a GEM:

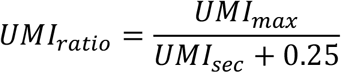

A small number (0.25) was added in the denominator to avoid division by zero.

The performance metric, o, is a weighted average of accuracy and fraction of retained GEMs, given by the following equation:

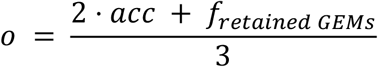

The accuracy metric is defined by the ratio of training set GEMs that were labeled as true interactions over the total number of GEMs in the training set. The performance metric, o, was maximized by finding the set of filters that increase the accuracy without excluding too much data.

The thresholds for filtering were selected from a complete grid search. Each feature was tested in the range of 0 to the median value, determined ad hoc from the experience that thresholds never approached the median value.

### Comparing TCR similarity

Effects of filtering were also evaluated through a comparison of TCR similarity. The similarity score is based on the kernel similarity score underlying the TCRmatch method between CDR3 sequences (Chronister et al., 2021). This score can be calculated for CDR3s of variable length, and takes a value between 0-1, with the value of 1 representing identical pairs. As both the α- and β-chain partake in the pMHC interaction, TCRs will be compared based on the summed similarity between the α- and β-chains, and GEMs missing a chain will be excluded to avoid bias. Two similarity scores are computed for each clonotype: an intra score and an inter score. The intra score is based on the maximum similarity of the given clonotype to all other clonotypes sharing its pMHC target (intra specificity). The inter score is based on the maximum similarity of the given clonotype to an equal sized set of clonotypes specific to other pMHC targets (inter specificity). The computation is done peptide-wise, such that clonotypes with maximum concordance for a given peptide will, for that peptide, be included in an intra similarity score, but for another peptide be included in an inter similarity score. Clonotypes consisting of GEMs causing diverging specificities were limited to the expected target pMHC or, if non-existing, to the specificity of highest concordance to avoid potential overlaps from “cross-reactive” clonotypes in the computation.

The similarity difference between intra and inter specificity clonotypes was tested for the hypothesis that intra similarity is greater than inter similarity (Wilcoxon, α=0.05). The similarity test was performed on all filtering methods described in the paper.

### Validating single-cell capture against fluorescent multimer staining responses

The 16 donors were known to respond to the panel of peptides used in the screening. Response proportions of sorted CD8+ T cells were detected by fluorescent multimer staining, as described previously. 1800 cells were selected from each donor and, based on the detected response proportions, an adjusted count of cells could be computed. Cells were selected based on two criteria: ≥10 cells and ≥ 0.002% of total CD8 T cells, or ≤10 cells but ≥0.01% of total CD8 T cells. The multimer responses were compared to GEMs filtered on UMI thresholds and matching HLA. To visually compare the two screening methods, the responses were normalized within each sample and plotted side-by-side. The methods were also quantitatively compared, both in absolute counts of responses and as binary classes with multimer responses as true labels and single-cell responses as query labels. The following evaluation metrics were computed based on binary classification of responses:

Matthew’s correlation coefficient (MCC)

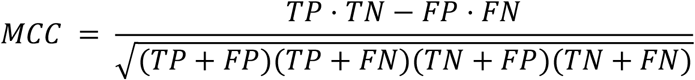

Recall (sensitivity or true positive rate)

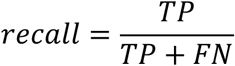

Precision (positive predictive value)

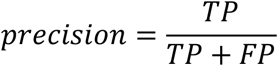

The following metrics were computed based on numerical values of responses, i.e. the adjusted number of cells and the count of GEMs.

Pearson correlation coefficient (PCC), where n is the number of screened peptides times the number of samples which have been screened. x and y represent the size of responses in single-cell screening and multimer staining, respectively.

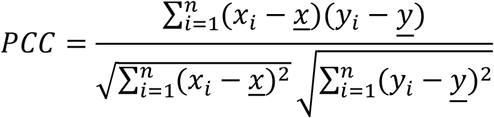

The correlation was also fitted via linear regression on log10 transformed data, resulting in the following equation.

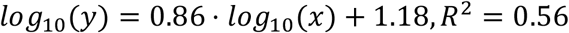

The equation was used to estimate the yield of single-cell captured cells relative to multimer screening. Three examples were computed to estimate an approximate 10% yield.

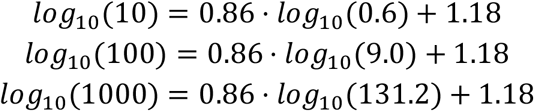

## Data availability

All data used for ATRAP filtering and analysis are available at https://github.com/mnielLab/ATRAP.

## Code availability

The ATRAP code is available at https://github.com/mnielLab/ATRAP.

## Supplementary information

### Supplementary tables

**Supplementary Table 1.** Samples. Overview of which samples contain cells from which donors and the relevant donor haplotypes.

| Hashing-ID | Donor | HLA-A | HLA-B |
| --- | --- | --- | --- |
| 1 | BC-300 | 0201 | 0702 |
| 2 | BC-326 | 0201 |  |
| 3 | BC-126 | 0201 |  |
| 4 | BC-328 | 0301 | 0702 |
| 5 | BC-62 | 0301 | 0702 |
| 6 | BC-355 | 0201 | 0702 |
| 7 | BC-360 |  | 0702 |
| 8 | BC-314 | 0301 | 0702 |
| 9 | BC-353 | 0301 | 0702 |
| 10 | BC-311, BC-11, BC-83, BC-88, BC-341, BC-342, BC-76 | 0201, 0301 | 0702 |

**Supplementary Table 2.**
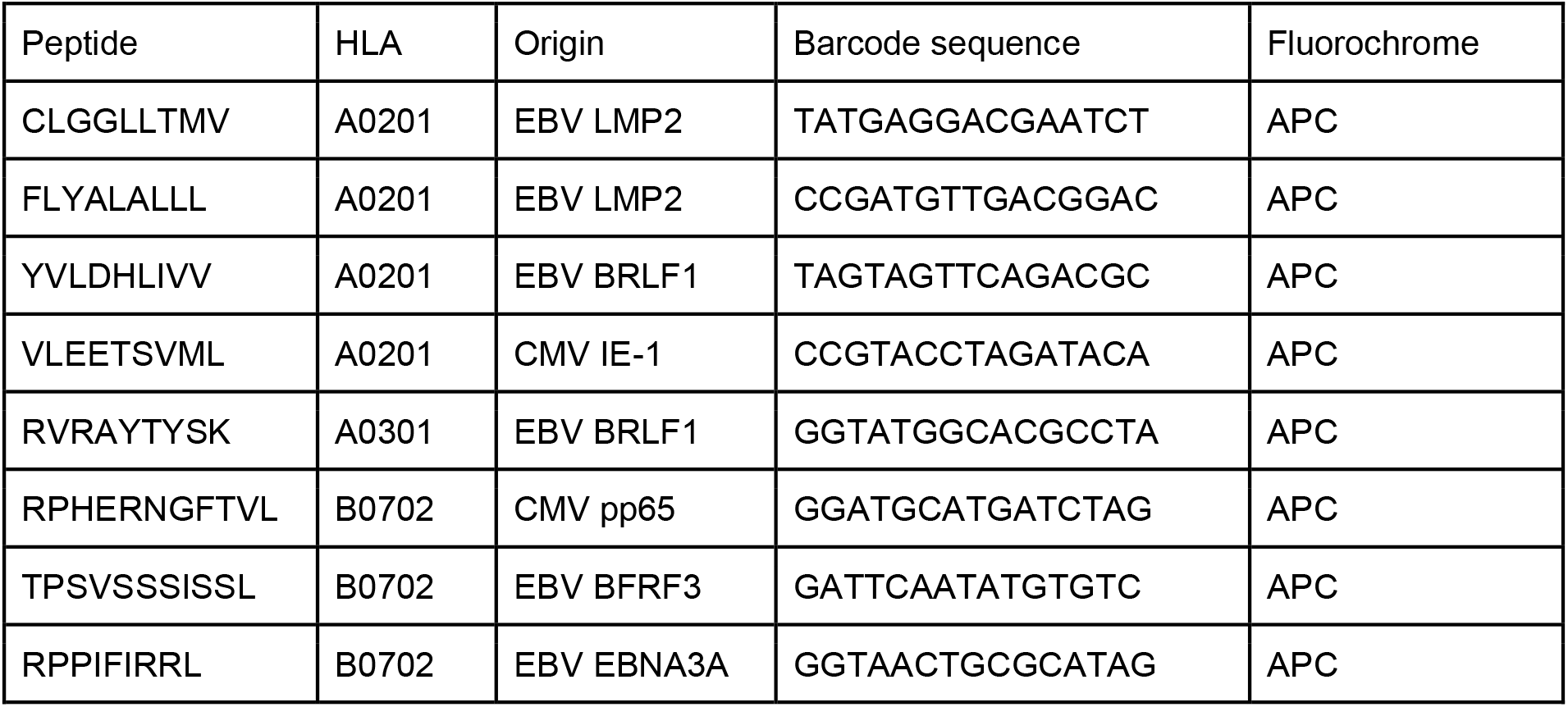
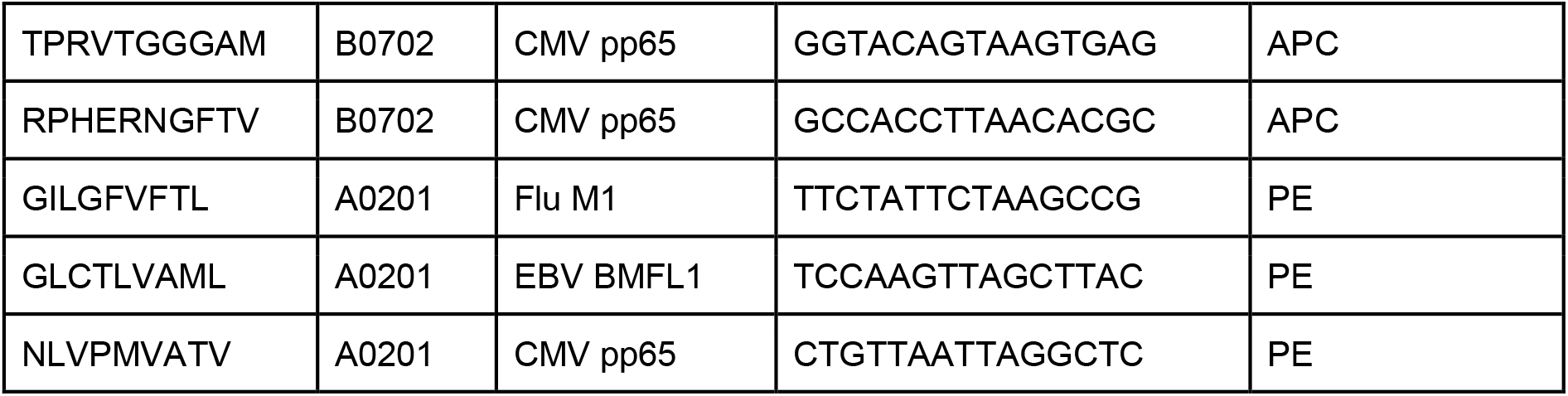
Peptide-MHC multimers. Information on the applied pMHC multimers. The full oligonucleotide tag are designed as follows: Biotin-C6-CGGAGATGTGTATAAGAGACAGNNNNNNNNNNXXXXXXXXXXXXXXXNNNNNNNNNCC CATATAAGAAA, with the barcode sequence indicated by 15 purple X’s. C6 indicates a six carbon spacer with a hydroxyl to the 5’ end of an oligonucleotide. Read2N is indicated by the black sequence, UMI’s are indicated in grey, and the capture oligo is indicated in turquoise.

**Supplementary Table 3.**
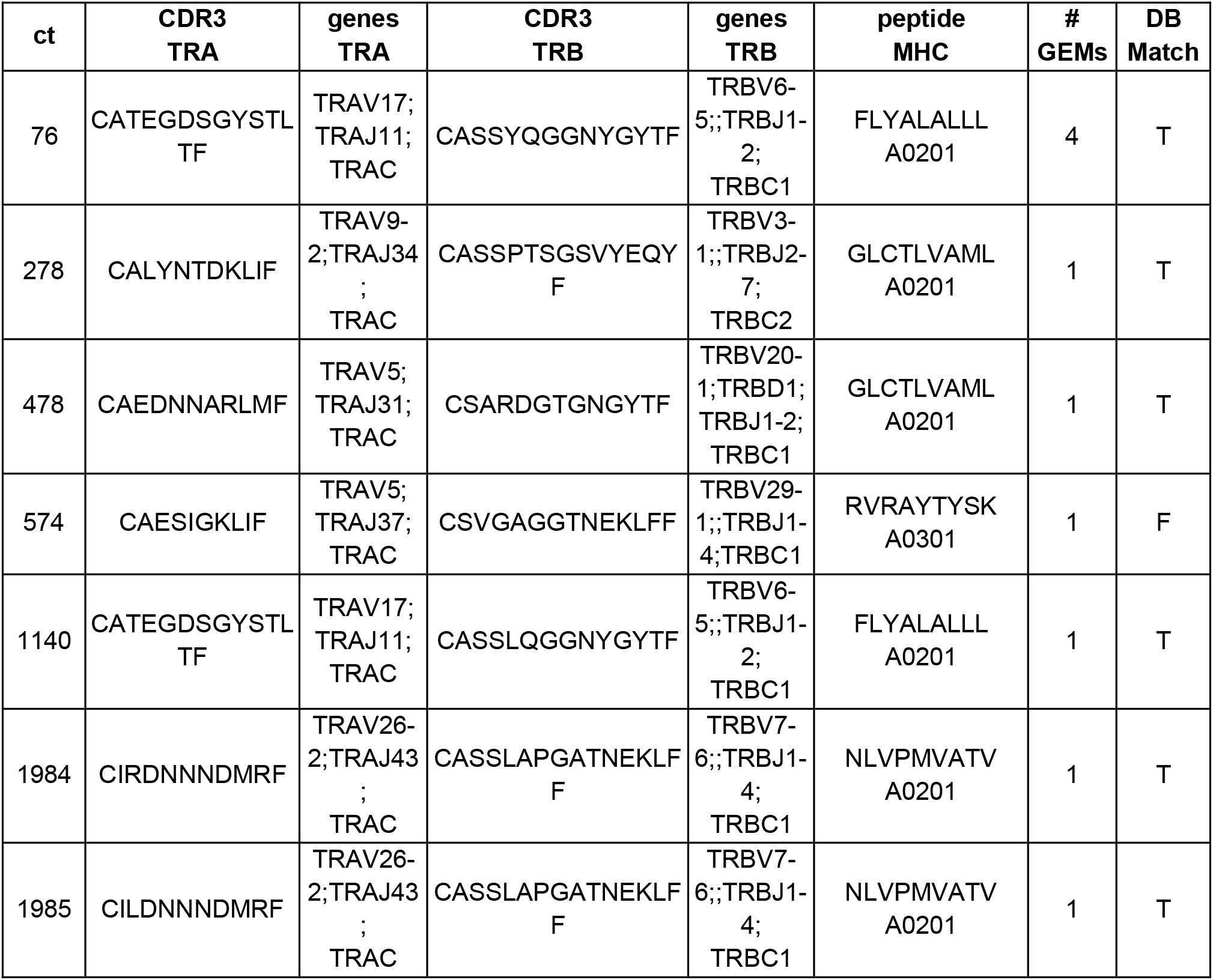
Database cross-referencing specificities. Information on the CDR3 sequences which matched the CDR3 sequences of the IEDB and VDJ databases presented in fig. 2d. Six different clonotypes (ct) had CDR3 sequence matches. Five of the clonotypes also matched (T:True) the database on the annotated pMHC (DB Match), while one clonotype (ct 573) had conflicting annotations.

**Supplementary table 4. Multimer staining responses**

All responses reported in Fig. 7. See table enclosed

## Supplementary figures

**Supplementary Figure 1:**
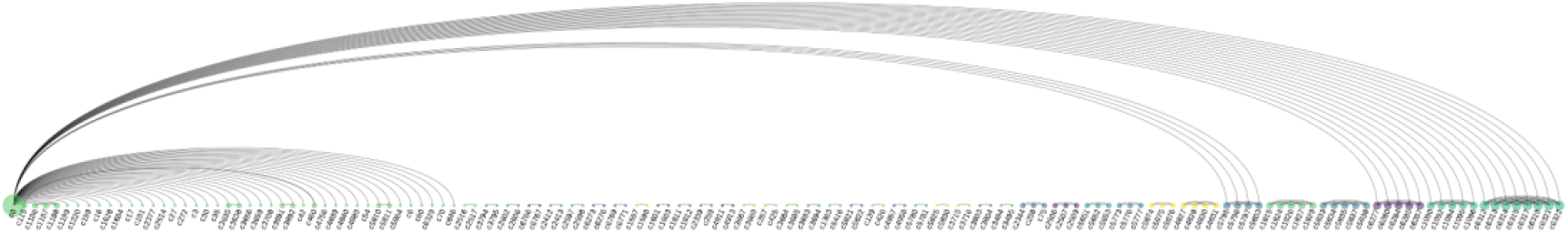
Clonotype replicas sharing VJ-CDR3ab. Arc diagram revealing shared VJab-genes and CDR3ab sequences across clonotypes defined by 10x Cellranger. Each node is a clonotype and the size reflects the magnitude GEMs in that clonotype sharing VJab-genes with GEMs of other clonotypes. The first node (c0, green) consists of the GEMs with no 10x clonotype annotation, while the remaining (c>0) are annotations by 10x Cellranger. The diagram reveals clonotype duplets (single arc connections), triplets (2 arcs), quadruplets, quintuplets, and even a single sextuplet. Since node c0 is a mixture of GEMs that were not annotated, the GEMs in this group will match many different clonotypes. Once a c0 VJ-CDR3 matches a clonotype which already is a replicate, the GEM will of course match all of them.

**Supplementary figure 2:**
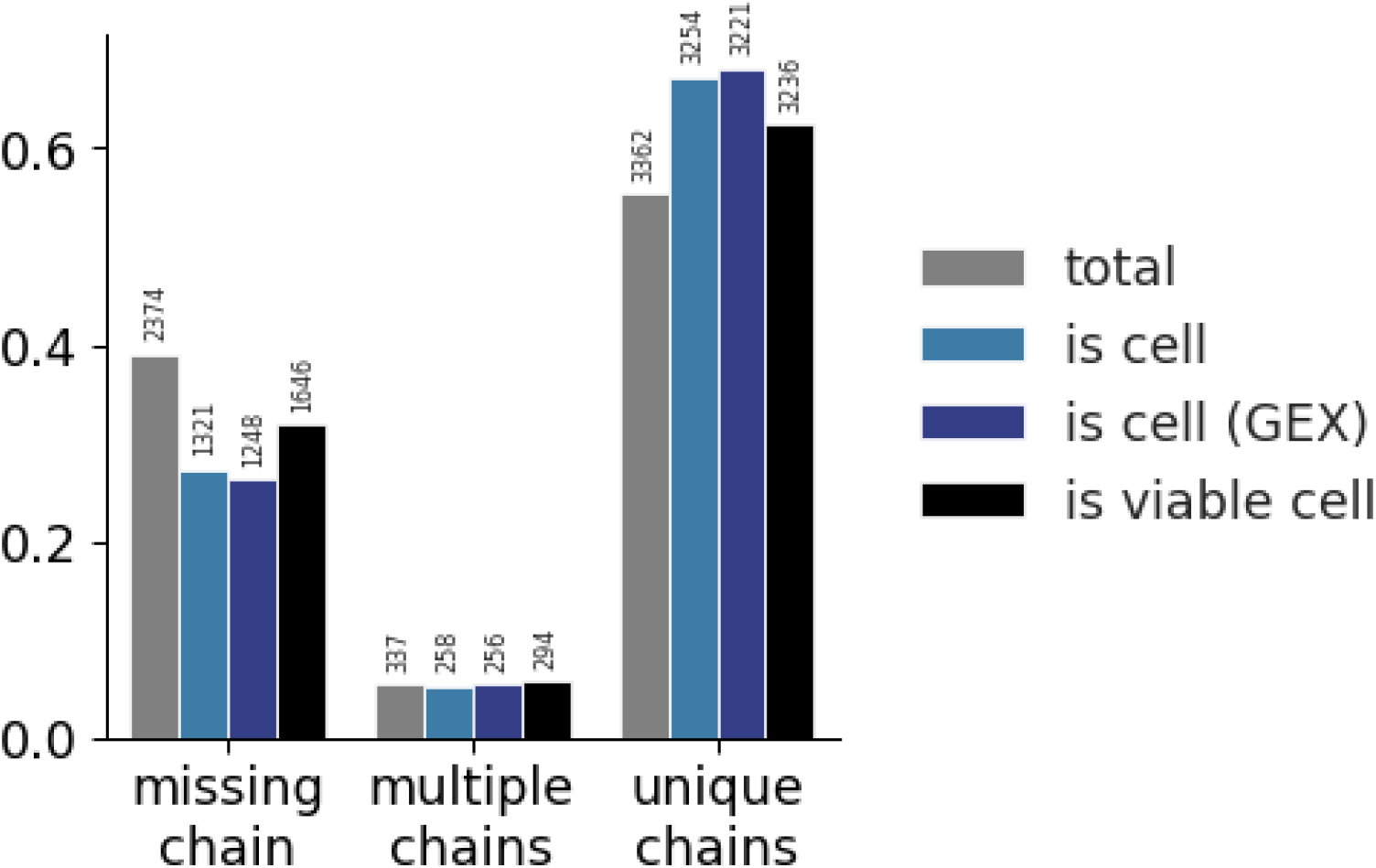
Distribution of the three categories of TCR chains across different methods of filtering. GEMs are categorized in one of three categories based on the detection of α- and β-chains: TCRs missing any chain, TCRs with multiple α- and/or β-chains, and TCRs with a unique set of one α- and one β-chain. The colors each represent a filtering step. The grey bars present the raw, total data with no filtering. The light blue bars present filtering on 10x Genomic’s Cellranger “is cell” call based on transcript level of TCR sequences only. The dark blue bars present filtering on 10x Genomic’s Cellranger “is cell” call based on transcript level of gene expression (GEX) sequencing. The black bars present filtering of GEX data on mitochondrial load and gene counts. For each step of filtering the counts within each category are normalized and the total value is listed above the bar. The raw data has a larger proportion of missing chain TCRs than the filtered sets. Filtering on “is cell” based on GEX data yields the largest proportion of unique chains. None of the filters completely nor substantially reduces the proportion of TCRs missing a chain or with multiple chains. See also supplementary note.

**Supplementary figure 3:**
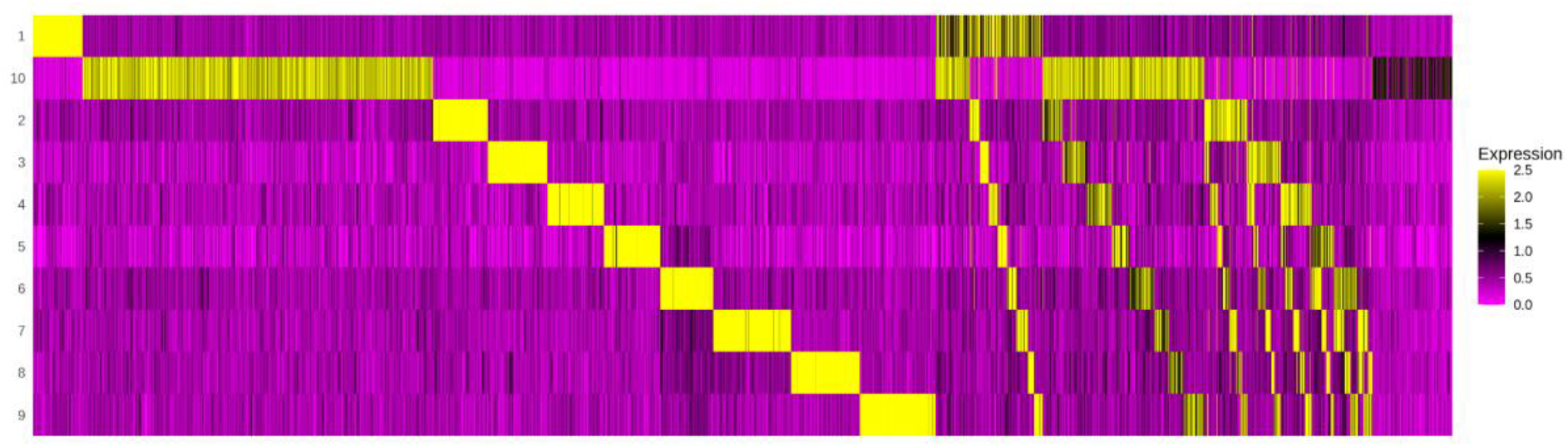
Demultiplexing cell hashing using Seurat. The GEMs (x-axis) are evaluated by the abundance of each sample barcode of 10 possible hashings (y-axis). The first section of the heatmap contains GEMs with unambiguous annotation to one sample. The second section illustrates how some GEMs contain barcodes for two samples, which might indicate a doublet, i.e. a capture of two T cells in one GEM. The last section reveals GEMs where no barcodes above a certain threshold were detected, and hence must be a result of leakage and can be discarded as noise.

**Supplementary figure 4:**
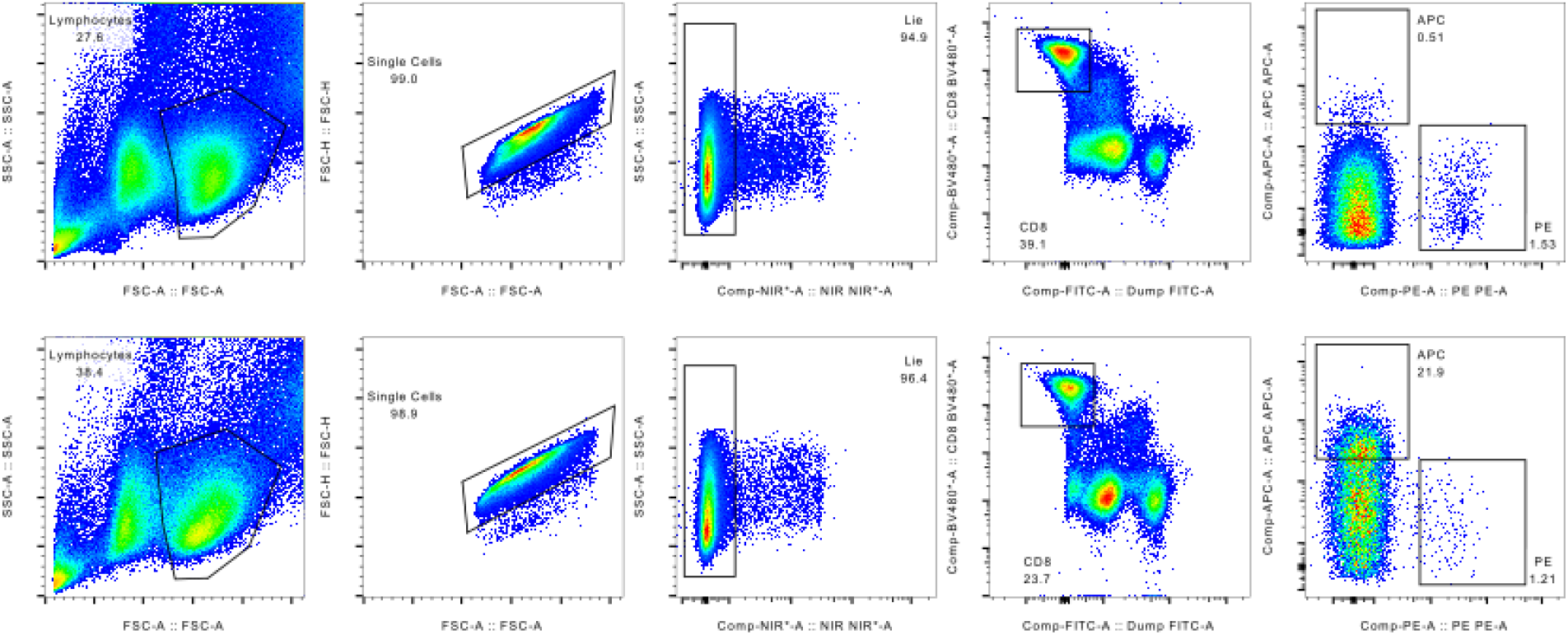
Gating strategy employed for sorting out pMHC binding MHC multimers isolated for single-cell processing.

## Supplementary note

### Additional filters to confidently assign the GEMs containing a cell

#### Removal of potential multiplets, leakage events, and dead cells

We set out to investigate how filters designed to remove potential multiplets, leakage events and dead cells would affect the distribution of GEMs between the three TCR categories: missing chain, multiple chains, and unique chains (Fig. 3). The 10x Genomics Software has a built-in method for flagging GEMs that are unlikely to contain a cell based on the transcript level of that GEM. Applying this filter based on the VDJ transcripts would reduce the set from 6073 to 4833. According to the software provider, the cell flagging method is more robust when including gene expression (GEX) data (10x Genomics 2022), which instead would reduce the set to 4725. Alternatively, the GEMs were filtered independently of Cellranger and directly on GEX data based on mitochondrial load and a minimum and maximum gene count per GEM, resulting in 5176 GEMs. The persisting GEMs should then be more likely to each contain a single viable T cell. Fig. 3 presents how filtering by the cell flag (‘is cell’) and viable cells affects the distribution of GEMs between the three TCR categories: missing chain, multiple chains, and unique chains. The filtered GEMs particularly contained TCRs which are missing an α- or a β-chain, however, the increased stringency of filtering did not substantially change the distribution of TCRs with unique chains relative to TCRs with missing or multiple chains. However, the filters substantially reduced the number of included GEMs (from 6073 without filters to 4725 when applying the most stringent filter).

such that the y-value of all three categories sum to 1. The distributions are shown for the unfiltered total GEMs (*total*), GEMs annotated as true cells by 10x Genomics Cellranger based on VDJ transcripts only (*is cell*), GEMs annotated as true cells when including GEX data (*is cell (GEX)*), and GEMs identified as viable cells from mitochondrial load and gene counts (*is viable cell*).

#### Applying hashing

The sample hashing component was predominantly observed as multiplets. In fact all, but one GEM, contained multiple sample hashing barcodes. An acknowledged method for demultiplexing SCseq data via sample hashing barcodes is the Seurat package: hashtag oligo (HTO) demultiplexing (Stoeckius et al. 2018). In short the method infers a threshold per sample barcode and thereby annotates GEMs as negative of any barcode, as singlets or doublets if multiple barcodes exceed their threshold.

Demultiplexing yielded 4,315 singlets, 1,580 doublets/multiplets, and 287 negatives (S.Fig. 9). Based on the large degree of ambient cell hashing barcode capture (Fig. 1a+b), we suspect that many of the 1,580 labeled doublets might be due to high contamination levels. Since demultiplexing was performed on the 6,073 GEMs containing both TCR and pMHC it is not surprising that only few GEMs are labeled negative.

In 320 GEMs, the demultiplexing method suggested another sample annotation than obtained from annotating by the most abundant barcode. Of the 320 GEMs, 283 were categorized as doublets and 37 singlets. The majority (227 & 35 GEMs) were originally annotated with sample 10. Since all three HLA alleles are contained in sample 10, any pMHC will inadvertently appear as having an HLA match. In only 8 GEMs the demultiplexing resulted in a different HLA profile, which corrected 7 GEMs from mismatches to matches between pMHC and sample HLA.

#### All responses reported in Fig. 7

The frequency or summed number of GEMs for all pMHCs reported in fig 7. The frequency found by fluorescent-based methods are utilized to calculate a proportion of each response per donor ((sum of responses in donor/100) x % of pMHC specific T cells) and then the estimated number of cells of each specificity sorted for the single-cell analyses ((total number of cells sorted x proportion)/100). In all cases a total of 1800 cells were sorted. Approximately 45% of sorted cells are lost already before loading on 10x, and 50% more are expected to be lost during 10x processing. Responses reported in italic are below

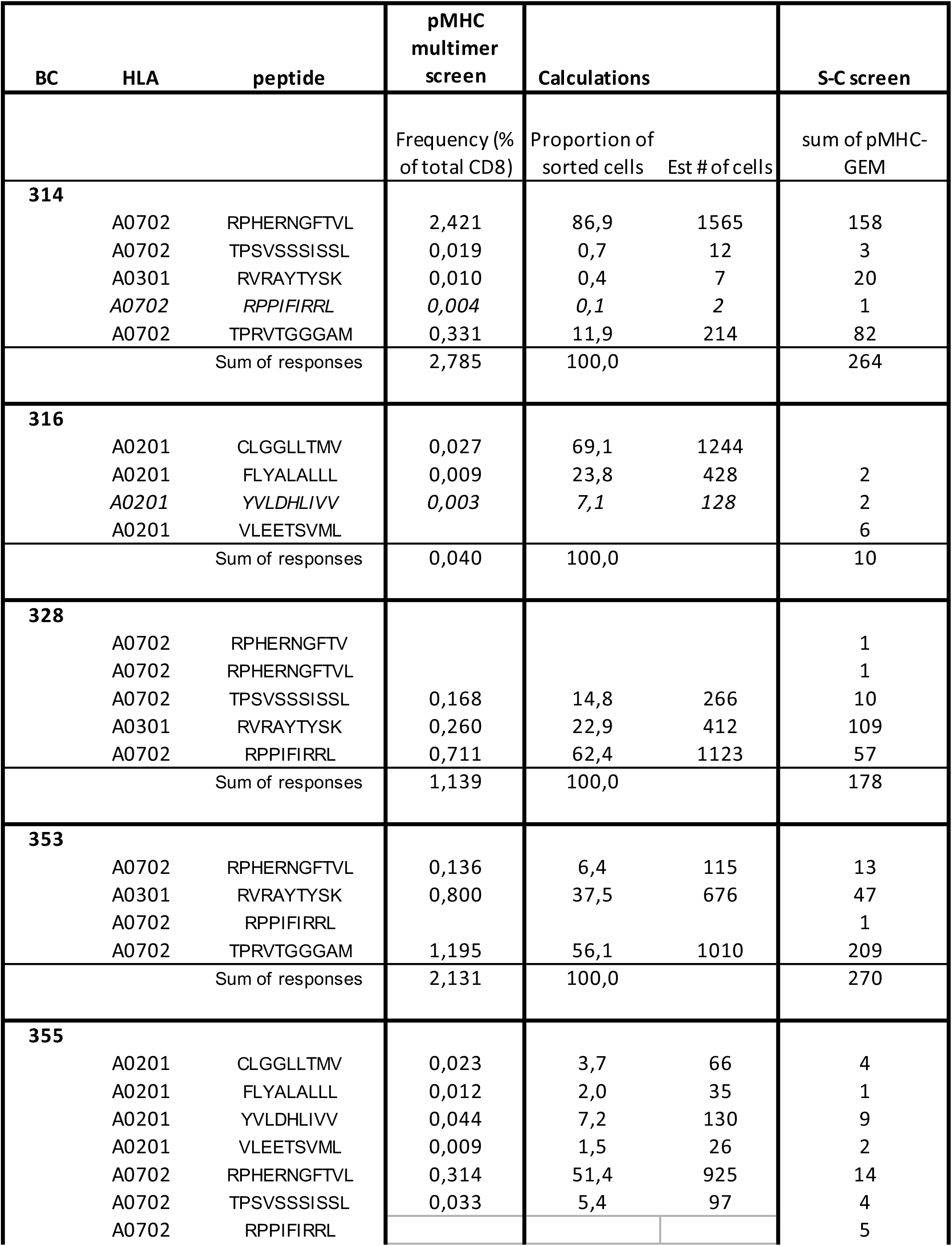
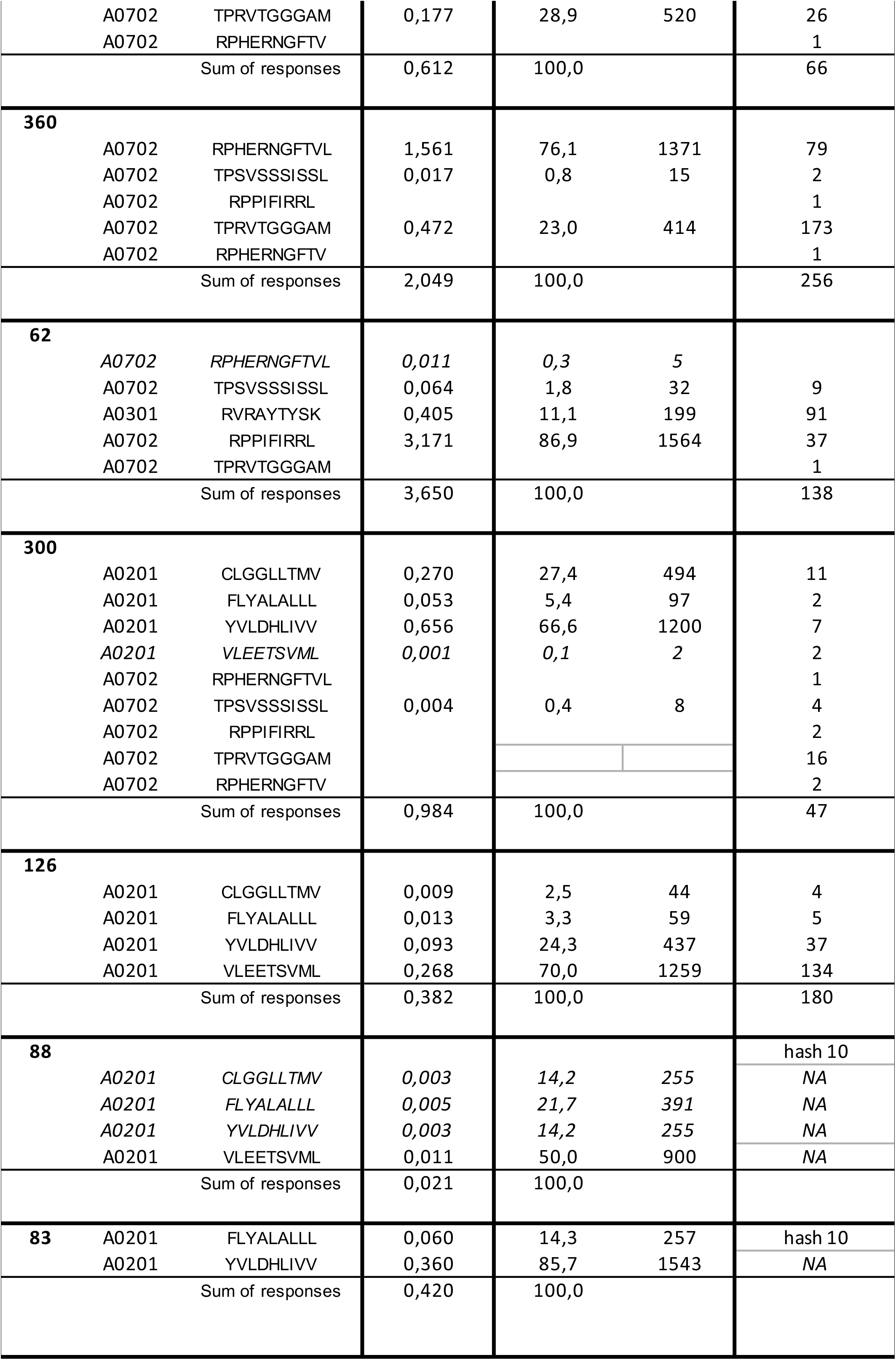
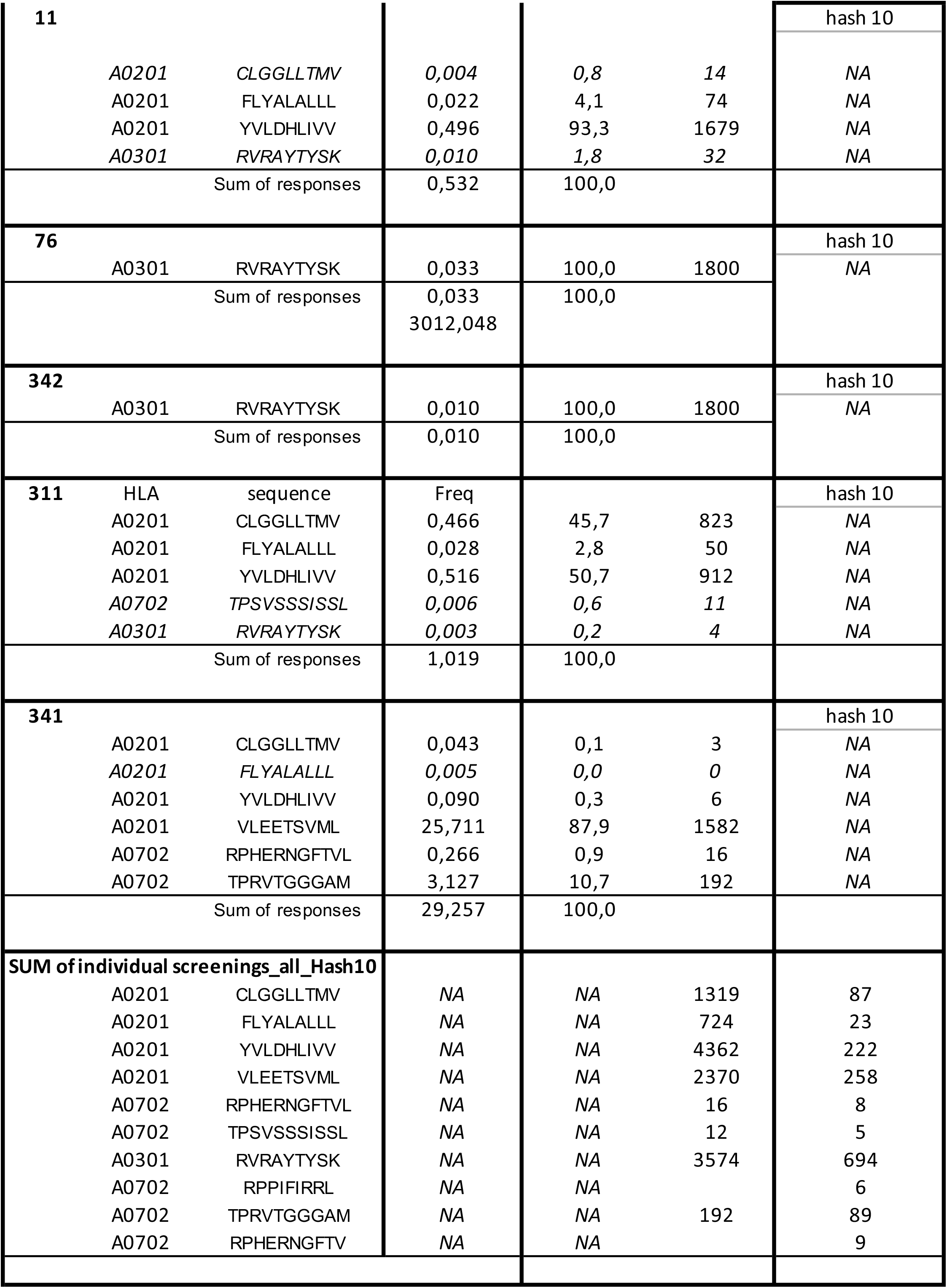

## References

10xGenomics. (n.d.-a). Cell Ranger Installation -Software -Single Cell Immune Profiling - Official 10x Genomics Support. Retrieved July 12, 2022, from https://support.10xgenomics.com/single-cell-vdj/software/pipelines/latest/installation

10xGenomics. (n.d.-b). V(D)J Cell Calling Algorithm -Software -Single Cell Immune Profiling -Official 10x Genomics Support. Retrieved July 12, 2022, from https://support.10xgenomics.com/single-cell-vdj/software/pipelines/latest/algorithms/cell-calling

Acha-Orbea, H., Mitchell, D. J., Timmermann, L., Wraith, D. C., Tausch, G. S., Waldor, M. K., … Steinman, L. (1988). Limited heterogeneity of T cell receptors from lymphocytes mediating autoimmune encephalomyelitis allows specific immune intervention. Cell, 54(2), 263–273. https://doi.org/10.1016/0092-8674(88)90558-2

Arstila, T. P., Casrouge, A., Baron, V., Even, J., Kanellopoulos, J., & Kourilsky, P. (1999). A direct estimate of the human alphabeta T cell receptor diversity. Science (New York, N.Y.), 286(5441), 958–961. https://doi.org/10.1126/SCIENCE.286.5441.958

Bagaev, D. V., Vroomans, R. M. A., Samir, J., Stervbo, U., Rius, C., Dolton, G., … Shugay, M. (2020). VDJdb in 2019: database extension, new analysis infrastructure and a T-cell receptor motif compendium. Nucleic Acids Research, 48(D1), D1057–D1062. https://doi.org/10.1093/NAR/GKZ874

Bakker, A. H., Hoppes, R., Linnemann, C., Toebes, M., Rodenko, B., Berkers, C. R., … Schumacher, T. N. M. (2008). Conditional MHC class I ligands and peptide exchange technology for the human MHC gene products HLA-A1, -A3, -A11, and -B7. Proceedings of the National Academy of Sciences of the United States of America, 105(10), 3825–3830. https://doi.org/10.1073/PNAS.0709717105/SUPPL_FILE/09717FIG7.JPG

Bentzen, A. K., Marquard, A. M., Lyngaa, R., Saini, S. K., Ramskov, S., Donia, M., … Hadrup, S. R. (2016). Large-scale detection of antigen-specific T cells using peptide-MHC-I multimers labeled with DNA barcodes. Nature Biotechnology 2016 34:10, 34(10), 1037–1045. https://doi.org/10.1038/NBT.3662

Bergman, R. (1999). How useful are T-cell receptor gene rearrangement studies as an adjunct to the histopathologic diagnosis of mycosis fungoides? The American Journal of Dermatopathology, 21(5), 498–502. https://doi.org/10.1097/00000372-199910000-00019

Bloom, J. D. (2018). Estimating the frequency of multiplets in single-cell RNA sequencing from cell-mixing experiments. PeerJ, 2018(9). https://doi.org/10.7717/PEERJ.5578/SUPP-4

Boutet, S. C., Walter, D., Stubbington, M. J. T., Pfeiffer, K. A., Lee, J. Y., Taylor, S. E. B., … Mikkelsen, T. S. (2019). Scalable and comprehensive characterization of antigen-specific CD8 T cells using multi-omics single cell analysis. The Journal of Immunology, 202(1 Supplement).

Buettner, F., Natarajan, K. N., Casale, F. P., Proserpio, V., Scialdone, A., Theis, F. J., … Stegle, O. (2015). Computational analysis of cell-to-cell heterogeneity in single-cell RNA-sequencing data reveals hidden subpopulations of cells. Nature Biotechnology 2014 33:2, 33(2), 155–160. https://doi.org/10.1038/nbt.3102

Chang, C. X. L., Tan, A. T., Or, M. Y., Toh, K. Y., Lim, P. Y., Chia, A. S. E., … Grotenbreg, G. M. (2013). Conditional ligands for Asian HLA variants facilitate the definition of CD8+ T-cell responses in acute and chronic viral diseases. European Journal of Immunology, 43(4), 1109–1120. https://doi.org/10.1002/EJI.201243088

Chronister, W. D., Crinklaw, A., Mahajan, S., Vita, R., Koşaloğlu-Yalçın, Z., Yan, Z., … Peters, B. (2021). TCRMatch: Predicting T-Cell Receptor Specificity Based on Sequence Similarity to Previously Characterized Receptors. Frontiers in Immunology, 12, 673. https://doi.org/10.3389/FIMMU.2021.640725/BIBTEX

Davis, M. M., & Bjorkman, P. J. (1988). T-cell antigen receptor genes and T-cell recognition. Nature, 334(6181), 395–402. https://doi.org/10.1038/334395A0

Dowell, A. C., Butler, M. S., Jinks, E., Tut, G., Lancaster, T., Sylla, P., … Ladhani, S. (2021). Children develop robust and sustained cross-reactive spike-specific immune responses to SARS-CoV-2 infection. Nature Immunology 2021 23:1, 23(1), 40–49. https://doi.org/10.1038/s41590-021-01089-8

Dziubianau, M., Hecht, J., Kuchenbecker, L., Sattler, A., Stervbo, U., Rödelsperger, C., … Babel, N. (2013). TCR Repertoire Analysis by Next Generation Sequencing Allows Complex Differential Diagnosis of T Cell–Related Pathology. American Journal of Transplantation, 13(11), 2842–2854. https://doi.org/10.1111/AJT.12431

Elliott, J. I., & Altmann, D. M. (1995). Dual T cell receptor alpha chain T cells in autoimmunity. The Journal of Experimental Medicine, 182(4), 953. https://doi.org/10.1084/JEM.182.4.953

Fluckiger, A., Daillère, R., Sassi, M., Sixt, B. S., Liu, P., Loos, F., … Zitvogel, L. (2020). Cross-reactivity between tumor MHC class I–restricted antigens and an enterococcal bacteriophage. Science, 369(6506), 936–942. https://doi.org/10.1126/SCIENCE.AAX0701/SUPPL_FILE/AAX0701_FLUCKIGER_SM.PDF

Frøsig, T. M., Yap, J., Seremet, T., Lyngaa, R., Svane, I. M., Thor Straten, P., … Hadrup, S. R. (2015). Design and validation of conditional ligands for HLA-B*08:01, HLA-B*15:01, HLA-B*35:01, and HLA-B*44:05. Cytometry Part A, 87(10), 967–975. https://doi.org/10.1002/CYTO.A.22689

Gaublomme, J. T., Li, B., McCabe, C., Knecht, A., Yang, Y., Drokhlyansky, E., … Regev, A. (2019). Nuclei multiplexing with barcoded antibodies for single-nucleus genomics. Nature Communications 2019 10:1, 10(1), 1–8. https://doi.org/10.1038/s41467-019-10756-2

Gerlach, C., Rohr, J. C., Perié, L., Van Rooij, N., Van Heijst, J. W. J., Velds, A., … Schumacher, T. N. M. (2013). Heterogeneous differentiation patterns of individual CD8+ T cells. Science (New York, N.Y.), 340(6132), 635–639. https://doi.org/10.1126/SCIENCE.1235487

Gielis, S., Moris, P., Bittremieux, W., De Neuter, N., Ogunjimi, B., Laukens, K., & Meysman, P. (2019). Detection of Enriched T Cell Epitope Specificity in Full T Cell Receptor Sequence Repertoires. Frontiers in Immunology, 10, 2820. https://doi.org/10.3389/FIMMU.2019.02820/BIBTEX

Hadrup, S. R., Toebes, M., Rodenko, B., Bakker, A. H., Egan, D. A., Ovaa, H., & Schumacher, T. N. M. (2009). High-throughput T-cell epitope discovery through MHC peptide exchange. Methods in Molecular Biology (Clifton, N.J.), 524, 383–405. https://doi.org/10.1007/978-1-59745-450-6_28

Hou, X., Wang, M., Lu, C., Xie, Q., Cui, G., Chen, J., … Diao, H. (2016). Analysis of the Repertoire Features of TCR Beta Chain CDR3 in Human by High-Throughput Sequencing. Cellular Physiology and Biochemistry : International Journal of Experimental Cellular Physiology, Biochemistry, and Pharmacology, 39(2), 651–667. https://doi.org/10.1159/000445656

Kharchenko, P. V., Silberstein, L., & Scadden, D. T. (2014). Bayesian approach to single-cell differential expression analysis. Nature Methods 2014 11:7, 11(7), 740–742. https://doi.org/10.1038/nmeth.2967

Kirsch, I. R., Watanabe, R., O’Malley, J. T., Williamson, D. W., Scott, L. L., Elco, C. P., … Clark, R. A. (2015). TCR sequencing facilitates diagnosis and identifies mature T cells as the cell of origin in CTCL. Science Translational Medicine, 7(308). https://doi.org/10.1126/SCITRANSLMED.AAA9122

Kivioja, T., Vähärautio, A., Karlsson, K., Bonke, M., Enge, M., Linnarsson, S., & Taipale, J. (2011). Counting absolute numbers of molecules using unique molecular identifiers. Nature Methods, 9(1), 72–74. https://doi.org/10.1038/NMETH.1778

Madi, A., Shifrut, E., Reich-Zeliger, S., Gal, H., Best, K., Ndifon, W., … Friedman, N. (2014). T-cell receptor repertoires share a restricted set of public and abundant CDR3 sequences that are associated with self-related immunity. Genome Research, 24(10), 1603–1612. https://doi.org/10.1101/GR.170753.113

Montemurro, A., Schuster, V., Povlsen, H. R., Bentzen, A. K., Jurtz, V., Chronister, W. D., … Nielsen, M. (2021). NetTCR-2.0 enables accurate prediction of TCR-peptide binding by using paired TCRα and β sequence data. Communications Biology, 4(1), 1–13. https://doi.org/10.1038/s42003-021-02610-3

Moris, P., De Pauw, J., Postovskaya, A., Gielis, S., De Neuter, N., Bittremieux, W., … Meysman, P. (2021). Current challenges for unseen-epitope TCR interaction prediction and a new perspective derived from image classification. Briefings in Bioinformatics, 22(4). https://doi.org/10.1093/BIB/BBAA318

Petrie, H. T., Livak, F., Schatz, D. G., Strasser, A., Crispe, I. N., & Shortman, K. (1993). Multiple rearrangements in T cell receptor alpha chain genes maximize the production of useful thymocytes. The Journal of Experimental Medicine, 178(2), 615. https://doi.org/10.1084/JEM.178.2.615

Robins, H. S., Campregher, P. V., Srivastava, S. K., Wacher, A., Turtle, C. J., Kahsai, O., … Carlson, C. S. (2009). Comprehensive assessment of T-cell receptor beta-chain diversity in alphabeta T cells. Blood, 114(19), 4099–4107. https://doi.org/10.1182/BLOOD-2009-04-217604

Rodenko, B., Toebes, M., Hadrup, S. R., van Esch, W. J. E., Molenaar, A. M., Schumacher, T. N. M., & Ovaa, H. (2006). Generation of peptide–MHC class I complexes through UV-mediated ligand exchange. Nature Protocols 2006 1:3, 1(3), 1120–1132. https://doi.org/10.1038/nprot.2006.121

Shen, W.-J., Wong, H.-S., Xiao, Q.-W., Guo, X., & Smale, S. (2012). Towards a Mathematical Foundation of Immunology and Amino Acid Chains. https://doi.org/10.48550/arxiv.1205.6031

Sherwood, J. (2013). Colonisation - it’s bad for your health: the context of Aboriginal health. Contemporary Nurse, 46(1), 28–40. https://doi.org/10.5172/CONU.2013.46.1.28

Sidhom, J. W., Larman, H. B., Pardoll, D. M., & Baras, A. S. (2021). DeepTCR is a deep learning framework for revealing sequence concepts within T-cell repertoires. Nature Communications 2021 12:1, 12(1), 1–12. https://doi.org/10.1038/s41467-021-21879-w

Stoeckius, M., Zheng, S., Houck-Loomis, B., Hao, S., Yeung, B. Z., Mauck, W. M., … Satija, R. (2018). Cell Hashing with barcoded antibodies enables multiplexing and doublet detection for single cell genomics. Genome Biology, 19(1), 1–12. https://doi.org/10.1186/S13059-018-1603-1/FIGURES/3

Toebes, M., Coccoris, M., Bins, A., Rodenko, B., Gomez, R., Nieuwkoop, N. J., … Schumacher, T. N. M. (2006). Design and use of conditional MHC class I ligands. Nature Medicine, 12(2), 246–251. https://doi.org/10.1038/NM1360

Vita, R., Mahajan, S., Overton, J. A., Dhanda, S. K., Martini, S., Cantrell, J. R., … Peters, B. (2019). The Immune Epitope Database (IEDB): 2018 update. Nucleic Acids Research, 47(D1), D339–D343. https://doi.org/10.1093/NAR/GKY1006

Weber, A., Born, J., & Rodriguez Martínez, M. (2021). TITAN: T-cell receptor specificity prediction with bimodal attention networks. Bioinformatics, 37(Suppl 1), i237. https://doi.org/10.1093/BIOINFORMATICS/BTAB294

Yamawaki, T. M., Lu, D. R., Ellwanger, D. C., Bhatt, D., Manzanillo, P., Arias, V., … Li, C. M. (2021). Systematic comparison of high-throughput single-cell RNA-seq methods for immune cell profiling. BMC Genomics, 22(1), 1–18. https://doi.org/10.1186/S12864-020-07358-4/FIGURES/8

Zhang, S. Q., Ma, K. Y., Schonnesen, A. A., Zhang, M., He, C., Sun, E., … Jiang, N. (2018). High-throughput determination of the antigen specificities of T cell receptors in single cells. Nature Biotechnology 2018 36:12, 36(12), 1156–1159. https://doi.org/10.1038/nbt.4282

Zhang, W., Hawkins, P. G., He, J., Gupta, N. T., Liu, J., Choonoo, G., … Atwal, G. S. (2021). A framework for highly multiplexed dextramer mapping and prediction of T cell receptor sequences to antigen specificity. Science Advances, 7(20). https://doi.org/10.1126/SCIADV.ABF5835

Zheng, G. X. Y., Terry, J. M., Belgrader, P., Ryvkin, P., Bent, Z. W., Wilson, R., … Bielas, J. H. (2017). Massively parallel digital transcriptional profiling of single cells. Nature Communications 2017 8:1, 8(1), 1–12. https://doi.org/10.1038/ncomms14049

